# D1 and D2 systems converge in the striatum to update goal-directed learning

**DOI:** 10.1101/780346

**Authors:** Miriam Matamales, Alice E. McGovern, Jia Dai Mi, Stuart B. Mazzone, Bernard W. Balleine, Jesus Bertran-Gonzalez

## Abstract

Extinction learning allows animals to withhold voluntary actions that are no longer related to reward and so provides a major source of behavioral control. Although such learning is thought to depend on dopamine signals in the striatum, the way the circuits mediating goal-directed control are reorganized during new learning remains unknown. Here, by mapping a dopamine-dependent transcriptional activation marker in large ensembles of striatal projection neurons (SPNs) expressing dopamine receptor type 1 (D1-SPNs) or 2 (D2-SPNs) in mice, we demonstrate an extensive and dynamic D2- to D1-SPN trans-modulation across the dorsal striatum that is necessary for updating previous goal-directed learning. Our findings suggest that D2-SPNs suppress the influence of outdated D1-SPN plasticity within functionally relevant striatal territories to reshape volitional action.

## Introduction

In changing environments, it is adaptive for humans and other animals to flexibly adjust their actions to maximize reward. One of the most powerful sources of behavioral change is extinction learning, which allows individuals to withhold instrumental actions when their consequences change. Rather than erasing such actions from one’s repertoire, current views propose that extinction generates new inhibitory learning that, when incorporated into previously acquired behavior, acts selectively to reduce instrumental performance (1).

Associative learning theory identifies the negative prediction errors produced by the absence of an anticipated reward as the source of the inhibitory learning underlying instrumental extinction (2). Such signals are thought to involve pauses in phasic dopamine (DA) activity, and this pattern is well suited to alter plasticity in the dorsomedial striatum (DMS), a key structure encoding the action-outcome associations necessary for goal-directed learning (3, 4). Nevertheless, the way multiphasic DA signals (5, 6) alter postsynaptic circuits in the DMS to shape goal-directed learning remains unknown.

Within the DMS, the plasticity associated with goal-directed learning involves glutamate release timed to local phasic DA activity to alter intracellular cyclic adenosine monophosphate (cAMP)-dependent pathways in postsynaptic neurons, a function that involves slow temporal scales (7) and that leads to gene transcription necessary for learning (8). This activity is distributed across two major subpopulations of spiny projection neuron (SPNs)—the principal targets of DA (9). These are completely intermixed within the striatum and express distinct DA receptor subtypes that respond to DA in an opposing manner (10): half express type 1 receptors and trigger powerful cAMP signaling in DA rich states (D1-SPNs), whereas the other half express type 2 receptors and show robust signaling in DA lean states (D2-SPNs) (11). Given that positive and negative prediction errors during appetitive learning are known to influence phasic DA release (12), we hypothesized that prediction errors during reward and extinction learning define distinctive molecular activation patterns in D1- and D2-SPN systems across the striatum to provide a molecular signature identifying those regions most relevant for plasticity.

Here we used wide-field, high-resolution imaging to reconstruct molecular signaling in D1- and D2-SPNs across whole striatal territories to assess how SPN subtypes encode both the excitatory and inhibitory learning that supports the acquisition and extinction of instrumental actions.

## Nucleosomal response in SPNs captures goal-directed learning

We first sought to establish whether intracellular signaling in SPNs undergoes functional reorganization across widespread territories of the striatum during goal-directed learning. We trained groups of mice to acquire rewarded instrumental actions, where a lever press (action) was associated with the delivery of a food pellet (outcome) (Fig. 1A and fig. S1A). In Group Early, initial acquisition of instrumental actions was marked by a spontaneous increase in lever press frequency on the first session of training, which was used to flag the approximate time at which the action-outcome contingency was first experienced (fig. S1, A to C). By contrast, mice in group Late received 12 days of additional instrumental training (fig. S1B) and developed well-established instrumental signatures, as revealed by an orderly increase in lever press responses measured at the center of the action-time space (fig. S1D).

**Fig. 1.**
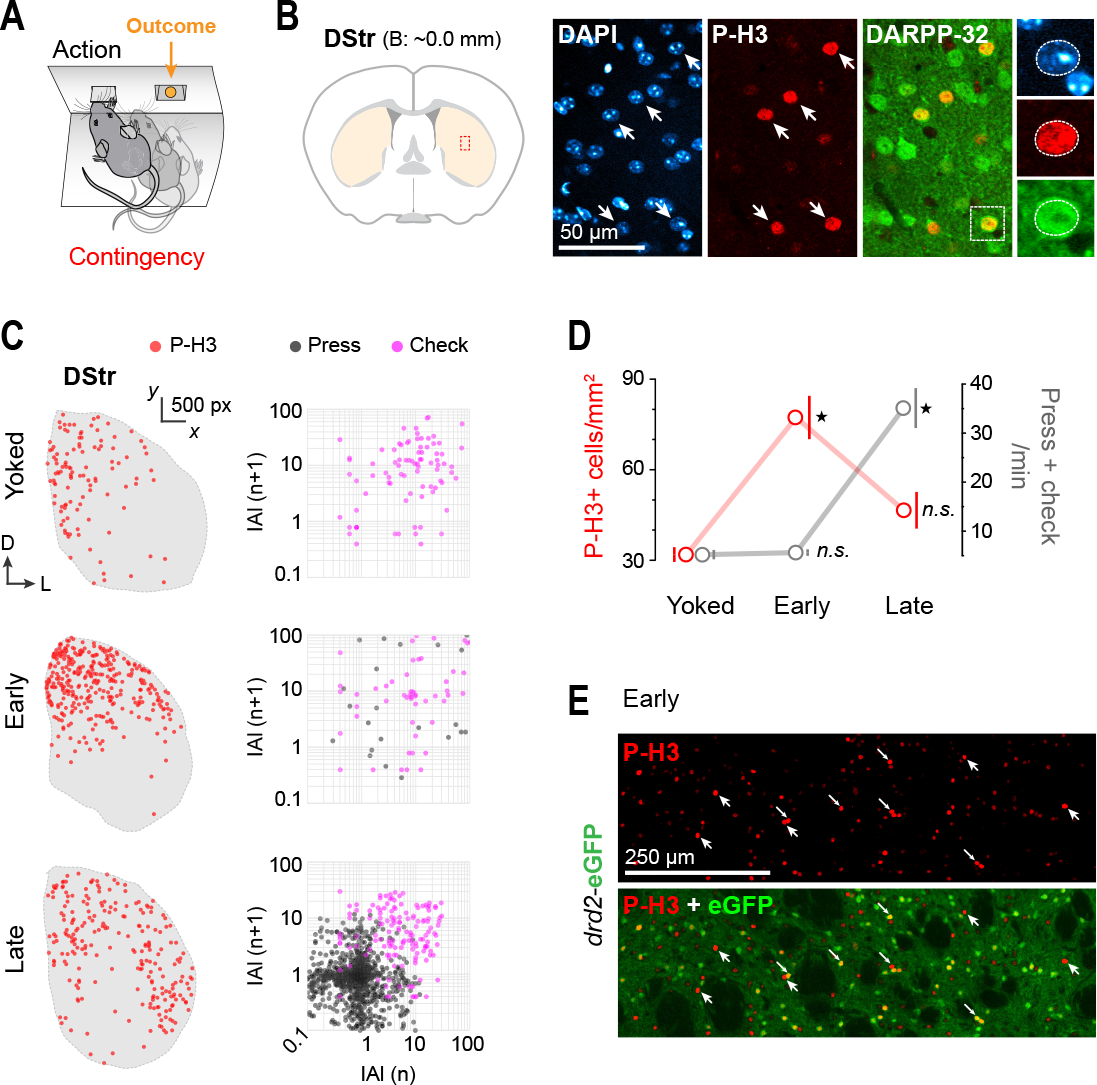
Nucleosomal response mapping reveals learning-related territories in the striatum. (**A**) Mice were trained to associate an action (lever press) with an outcome (food pellet) (fig. S1). (**B**) Immunodetection of phosphorylated nucleosomes (phospho-Ser10-histone H3; P-H3) identifies transcriptionally active (ta) neurons in the dorsal striatum (DStr). P-H3 immunoreactivity was specifically detected in the nuclei (DAPI+) of SPNs (DARPP-32+). (**C**) Digitized reconstruction of taSPNs throughout the DStr in “Early”, “Late” and “Yoked” mice (7,455 SPNs mapped). Right panels: return maps of inter-action-intervals (IAIs) for lever presses (grey) and magazine checks (purple). (**D**) taSPN density (left axis, red) and overall action rate (right axis, grey) in the different training groups. (**E**) Identification of taSPNs in the DStr of a trained drd2-eGFP mouse. Arrowheads and arrows: taD2- and taD1-SPNs, respectively. *, simple effects. N.S., non-significant (Table S1).

We next assessed whether the different training received by these mice was captured in the signaling patterns detected in striatal SPNs. We used immunodetection of the phosphorylated form of histone H3 on serine 10 (P-H3), a ubiquitous transcriptional activation marker that is rapidly induced in SPNs in response to different DA states (8, 10, 13, 14). We found a robust P-H3 signal in the nucleus of striatal neurons that co-labeled with DARPP-32, a marker of SPNs (Fig. 1B), suggesting that projection neurons—relative to other types of striatal neuron—were transcriptionally active under these conditions. Wide-field, high-resolution mapping identified significantly different levels of transcriptionally active SPNs (taSPNs) across groups, with clear territorial differences in their distribution (Fig. 1, C and D). Compared to Yoked controls, that were exposed to the lever and received as many rewards but non-contingently, the Early group showed a high density of taSPNs concentrated in the DMS (Fig. 1C), consistent with the role of this region in action-outcome encoding (4). In contrast, group Late showed a significantly reduced and sparsely distributed population of taSPNs that extended to the dorsolateral striatum (DLS; Fig. 1C), in support of the functional lateralization expected from extensively trained actions (15, 16). Critically, we found a clear dissociation between taSPN density and the extent of overall performance (i.e. lever presses and magazine checks) (Fig. 1, C and D), confirming that our measures of neuronal activation reflected learning rather than movement execution in the striatum (Fig. 1D; table S1). This allowed us to link goal-directed learning with the induction of DA-promoted transcriptional activity in postsynaptic SPNs. Importantly, the nuclear P-H3 signal was detected in D1- as well as D2-SPN subtypes (Fig. 1E), indicating that both of these neuronal systems were responsive to the fluctuating DA states underpinning goal-directed learning.

## Regional overlap of activated SPN subpopulations predicts extinction learning

To study the activation patterns of D2- and D1-postsynaptic circuits in the striatum during instrumental learning and extinction, we mapped and classified large numbers of taSPNs in whole striatal sections of drd2-eGFP mice (fig. S2). We trained two groups of mice on an increasing fixed ratio (FR) reinforcement schedule where access to each food outcome relied on a predictable and well-defined instrumental effort (Fig. 2A). The groups showed indistinguishable performance with very similar elevation of lever press rate across training (fig. S3A, table S2). On day 16, one group (Extinction) underwent an altered training session in which lever pressing activated the food dispenser, but no outcomes were delivered. This manipulation generated vigorous responding for “no-reward” (Ø) that was comparable to that of non-extinguished mice (Instrumental controls) for almost half of the session, at which point their cumulative performance decayed (Fig. 2B; fig. S3B; table S2).

**Fig. 2.**
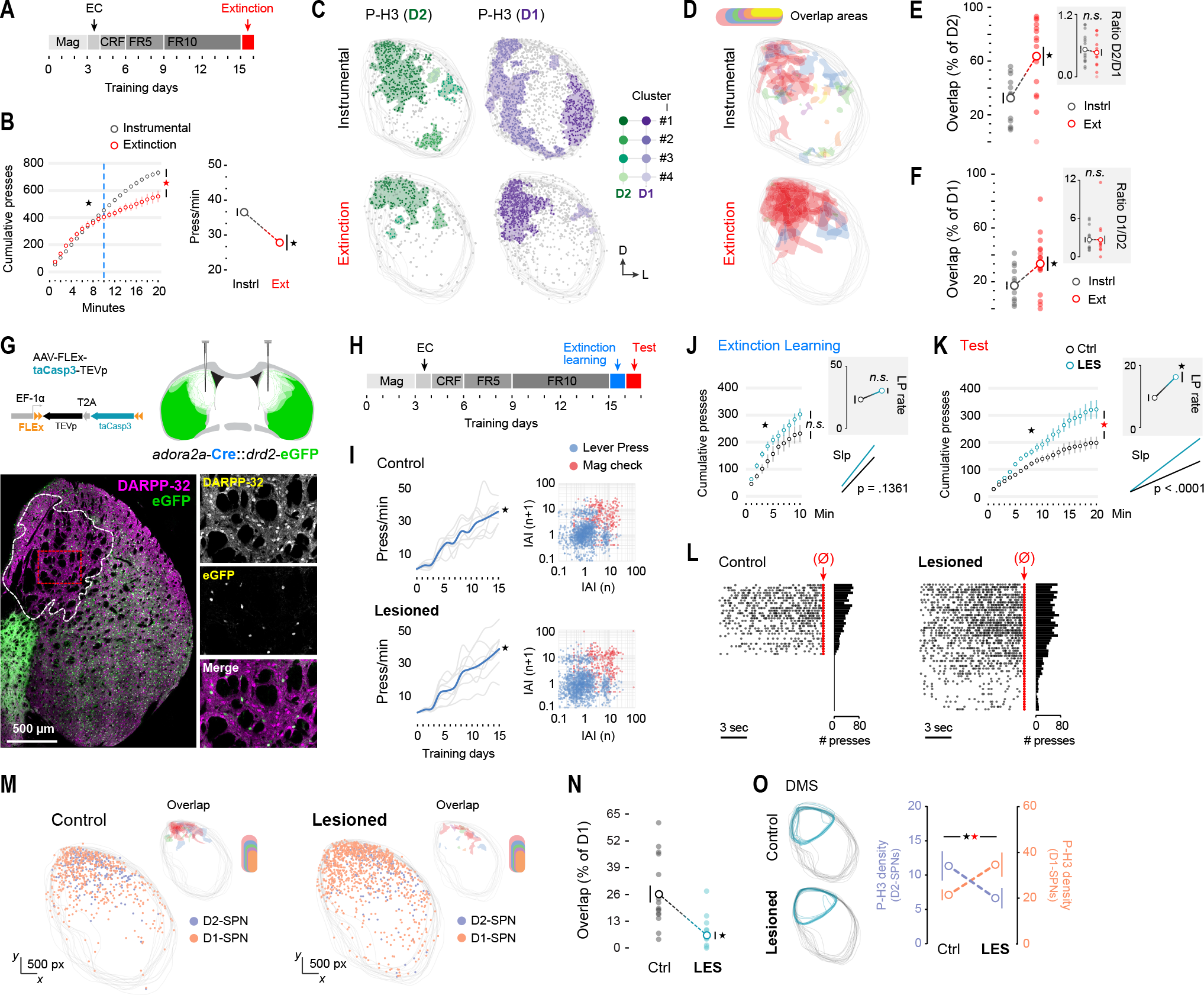
Functional confluence of projection systems in the striatum promotes extinction of learned actions. (**A**) Mice were trained to increasing fixed ratio (FR) reinforcement schedules prior to extinction (day 16). (**B**) Cumulative and average lever press performance during instrumental (control) and extinction sessions (day 16). (**C**) Distribution taD2- and taD1-SPNs in the dorsal striatum (DStr) of mice in (B). Plots show up to 4 density clusters from aligned hemisections in each group (4,044 SPNs mapped). (**D**) Reconstruction of taD2- and taD1-SPN overlapping territories in the DStr. Color code: size of overlap areas in each image. (**E-F**) Extent of taD2- and taD1-SPN territory overlap. Data are % of taD2-SPN territories occupied by taD1-SPNs (E) and vice versa (F). Insets: overall D2/D1 (E) and D1/D2 (F) P-H3+ SPN ratios. (**G**) Genetic lesion of D2-SPNs in the dorsomedial striatum (DMS) through AAV-FLEx-ta-Casp3-TEVp system. DARPP-32+-eGFP-SPNs remained intact (fig. S4A). (**H**) Control and Lesioned mice were trained as in A, but underwent additional extinction testing on day 17. (**I**) Lever press performance in both groups across instrumental training. Right: return maps of collective inter-action-intervals (IAIs) on day 15. (**J-K**) Cumulative presses/min on days 16 (J) and 17 (K). Insets: average press performance (top) and linear regression slope (Slp) analysis (bottom). (**L**) Raster plots and frequency histograms of pooled lever press data preceding the delivery of each pseudoreward (red). (**M**) Digitized reconstruction of taD2- and taD1-SPNs after test. Insets: taD2- and taD1-SPN overlapping territories. (**N**) Extent of taD2- and taD1-SPN territory overlap (% of D1). (**O**) P-H3+ nuclei density in D2-SPNs (left axis, purple) and D1-SPNs (right axis, orange) in the DMS. Left: regions quantified in each group. *, overall/simple effect (black) and interaction (red). N.S., non-significant (Table S2).

Mapping of taSPNs in entire striatal sections revealed that overall densities of P-H3+ D2- and D1-SPNs were similar in Instrumental and Extinction groups (fig. S3 C and D, table S2). However, pooled cluster analysis, conducted on each system separately, showed that taD2- and taD1-SPNs followed characteristic spatial distributions across the dorsal striatum in each group (Fig. 2C). We found that, in mice undergoing a rewarded session, each system colonized alternative areas in the DMS, and taD1-SPNs alone segregated to exclusive DLS territories (Fig. 2C, top). By contrast, in animals undergoing extinction, we found a high level of convergence in DMS areas, and very few neurons were detected laterally (Fig. 2C, bottom). Direct assessment of the overlap between clusters of taD2- and taD1-SPNs in each individual section confirmed that Extinction mice exhibited a marked increase in the proportion of taD2-SPN territories that coincided with functional D1-SPN areas specifically in the DMS (Fig. 2 D and E), as well as a higher proportion of taD1-SPN clusters sharing space with taD2-SPNs in this same region (Fig. 2D and F) (table S2). Our results show, therefore, that areas with highly convergent D2- and D1-SPN activation can define regions with elevated plasticity in the dorsal striatum and suggest that striatal projection systems dynamically interact with one another to establish functional territories during learning.

## D2-SPNs in the DMS are required to encode extinction learning

We hypothesized that recruiting activated D2-SPNs in the DMS is directly related to inhibitory learning during extinction. We addressed this specific claim by selectively removing D2-SPNs from the DMS where we had observed high taD2- and taD1-SPN confluence (see Fig. 2D). We used the taCasp3-TEVp genetic lesion system (17) to induce robust and selective ablations of D2-SPNs in the DMS in adult adora2a-Cre∷drd2-eGFP hybrid mice (Fig. 2G and fig. S4A). After recovery, Lesioned and Control mice were given instrumental training, as previously. On day 16, they were given a 10-min extinction learning session and then, 24h later (day 17), received an extinction test (Fig. 2H). This protocol was aimed at detecting differences on test due to deficient integration of extinction learning 24 hours earlier. Importantly, D2-SPN ablation had no effect on the initial acquisition of goal-directed action; Control and Lesioned groups showed very similar levels of performance across sessions (Fig. 2I, left) and an indistinguishable response structure on the last day of training (Fig. 2I, right) (table S2). This suggested that D2-SPNs may not be directly involved in the initial acquisition of goal-directed action, as proposed previously (18). Likewise, both Lesioned and Control groups similarly reduced lever press performance during the 10-min extinction learning session (Fig. 2J), indicating that D2-SPN ablation did not affect performance during initial extinction (table S2). By contrast, performance during the test session 24h later differed significantly: Lesioned mice accumulated a higher number of lever presses across the session and showed a higher average level of lever pressing as well as a steeper linear regression slope (Fig. 2K; table S2). This increase in performance was not due to an overall increase in lever press rates but rather recurrent and persistent performance during the extinction session (Fig. 2L; fig S4, B to D; table S2).

We then analyzed the distribution of taD2- and taD1-SPNs that accumulated in the dorsal striatum during the 20-min test on day 17. Consistent with our previous result, we again found an elevated confluence of taD2- and taD1-SPNs in the DMS of control mice, which again displayed a significant overlap despite being assessed on the second day of extinction (Fig. 2, M and N; table S2). Conversely, the absence of D2-SPNs in Lesioned mice was associated with a strikingly high density of taD1-SPNs in the DMS, suggesting that molecular activity in D1-SPNs was released from modulatory control (Fig. 2, M and N; table S2). Density analysis in the DMS revealed that taD2- and taD1-SPNs followed opposing density patterns in Control and Lesioned mice: higher densities of taD2-SPNs predicted low densities of taD1-SPNs and vice versa (Fig. 2O; table S2). These results suggest, therefore, that the functional recruitment of D2-SPNs in the DMS is required to modulate plasticity in neighboring D1-SPNs during extinction learning.

## D2-SPNs spatially rearrange D1-SPN plasticity

Given this conclusion, we sought to establish whether D2- and D1-SPNs functionally interact within convergent striatal territories using pharmacological compounds known to induce robust and widespread intracellular signaling in each system (13, 19). Systemic injection of raclopride (RAC; a DA D2 receptor antagonist) to drd2-eGFP mice induced a strong nucleosomal response mostly in D2-SPNs that extended throughout the dorsal striatum, whereas GBR12783 (GBR; a DA transporter inhibitor that elevates extracellular DA) induced an even stronger activation with similar distribution but mostly in D1-SPNs (Fig. 3A). We injected 4 groups of drd2-eGFP mice with different combinations of vehicle, RAC (t = 0) and GBR (t = 15) and recorded their ambulatory locomotor activity in an open field arena prior to perfusion (t = 30) (Fig. 3B). We found that GBR injection strongly increased locomotion in both groups receiving it, irrespective of the RAC injection (Fig. 3, C to G, table S3). In the striatum, we found opposing patterns of taD2- and taD1-SPN clusters: in RAC-treated mice taD2-SPNs dominated most of the striatal space (Fig. 3, H and I), whereas a reverse pattern was observed after GBR treatment (Fig. 3,). The combination of both drugs, however, resulted in a high density of taD2-SPNs and a low density of taD1-SPNs throughout the dorsal striatum, indicating that prior injection of RAC prevented the effects of GBR on D1-SPN plasticity (Fig. 3K). Moreover, the distribution of taD2 and taD1-SPN peak density areas tended to occupy alternative spaces in the dorsal striatum in all groups, regardless of the pharmacological cocktail received (Fig. 3, H to K, lower panels). This opposition was supported by a strong overall treatment × neuron interaction (Fig. 3L; table S3). Paradoxically, animals with D2-dominated SPN plasticity (Fig. 3K) showed unaltered GBR-induced hyperlocomotion (Fig. 3G), again suggesting that nuclear plasticity in SPNs can be dissociated from the expression of behavioral performance (cf. Fig. 1D). These results provided further evidence that, in territories where both systems functionally converge, taD2-SPNs directly modulate D1-SPN plasticity.

**Fig. 3.**
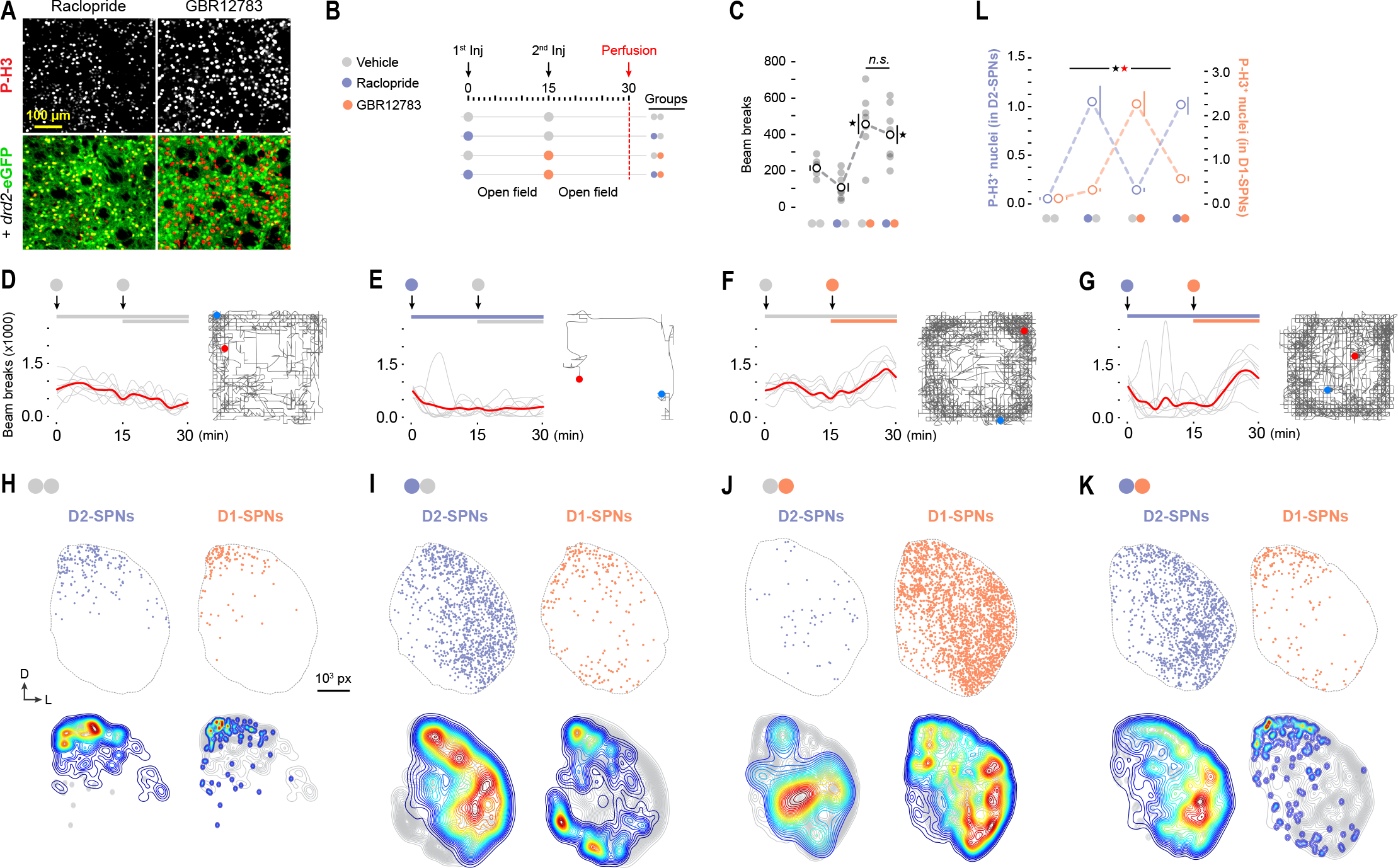
Overstimulated SPN systems compete for space in the striatum. (**A**) Confocal micrographs showing the effects of D2R blockade (Raclopride, 0.3 mg/kg) and DAT inhibition (GBR12783, 15 mg/kg) on taSPNs. (**B**) Two-injection experimental design applied to each group prior to perfusion. Ambulation was measured in an open field. (**C**) Ambulation (beam brakes) recorded after the second injection (min 15-30). (**D-G**) Ambulatory activity per min. Right: ambulatory trajectory (start: blue; finish: red) in one example mouse after the second injection. (**H-K**) Top: maps of taD2- and taD1-SPNs in the dorsal striatum (DStr) of an example mouse. Bottom: distribution contour plots delimitating regions of increasing 3+ nuclear density in D2- (left) and D1- (right) SPN systems separately. Isodensity curves are pseudocolored from low (blue) to high (red) relative densities (31,542 taD1-SPNs and 29,100 taD2-SPNs mapped). (**L**) Quantification of P-H3+ nuclei (counts x 103) distributed in D2- and D1-SPNs in each pharmacological condition. *, significant overall/simple effect (black) and interaction (red). N.S., not significant (Table S3).

## Broad connectivity of the local striatal network supports an extensive D2-to-D1-SPN trans-modulation

Based on these findings, we reasoned that the local network in the dorsal striatum may reflect connectivity biases consistent with a direct D2-to-D1 modulation, in line with that proposed in other striatal areas (20–23). In order to verify that both SPN types produce local terminals in the striatum, we labelled axonal projections and synaptic boutons in D1 and D2 systems separately using Cre-dependent expression of membrane-bound GFP (mGFP) and synaptophysin-driven mRuby (Syp-mRuby) (24) (fig. S5, A to C). In the striatum, both SPN subtypes were shown to provide abundant axonal terminals proximal to the injection site (fig. S5D) organized around distal dendrites (insets), proximal dendrites, somatic regions and virtually all other histological compartments, consistent with the proposed heterogeneous nature of SPN collaterals (25).

We next sought to investigate the overall magnitude of the connectivity between SPN types in large areas of the striatum. We used a quantitative network-level approach to measure connectivity biases using unilateral injections of the herpes simplex (HSV1) H129tdTomato virus in the dorsal striatum of drd2-eGFP mice (Fig. 4A). Since this virus moves along synaptically connected neurons in the anterograde direction (26), we mapped transduced tdTomato+ SPNs across large territories to reveal the broad connectivity patterns established within the striatum (Fig. 4B, fig. S6A). We found that only ~20% of the transduced SPNs were D2-SPNs, whereas up to ~80% were D1-SPNs (Fig. 4, C to E; table S4). These numbers were remarkably similar across all areas of infection, regardless of whether they were ipsi- or contralateral to the initial injection (fig. S6, B to D; table S4). Moreover, an injection of the same virus in the prelimbic cortex (i.e. one upstream synapse) provided very similar weights of transduced D1- (~80%) and D2- (~20%) SPNs (fig. S6, E to G; table S4), suggesting that the overall connectivity was substantially biased towards D1-SPNs, and that this pattern was established locally within the striatum.

**Fig. 4.**
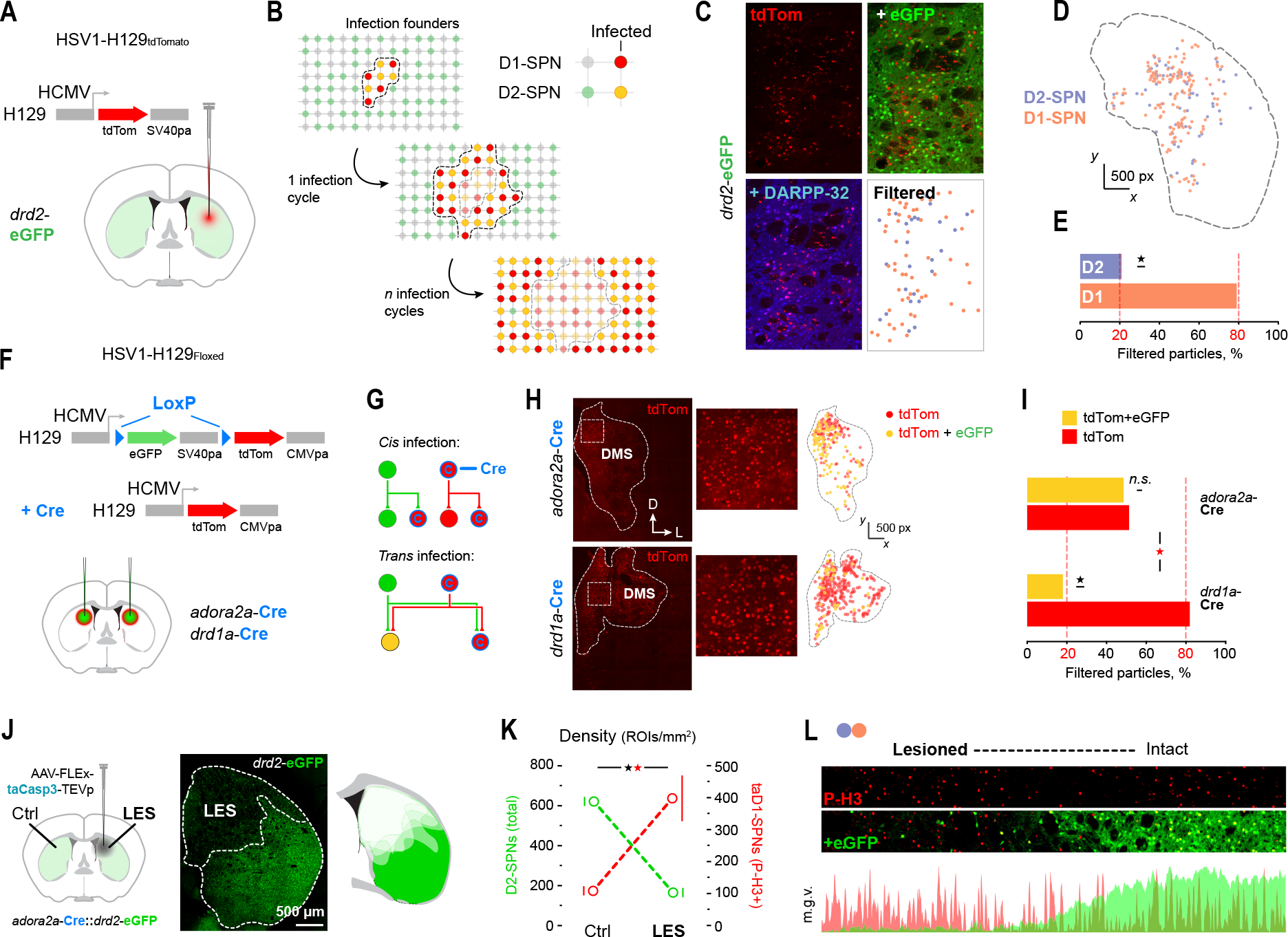
SPN subtypes show large-scale unidirectional trans-connectivity. (**A**) Anterograde trans-synaptic HSV1-H129-td-Tomato virus was unilaterally injected in the dorsal striatum (DStr) of drd2-eGFP mice. (**B**) In a hypothetical striatum with symmetrical SPN-SPN contacts, the “founder” cells first integrating the virus are expected to infect similar proportions of surrounding SPNs (D2 and D1). (**C**) Infected, Td-Tomato+, particles were classified as D2- and D1-SPNs according to their eGFP and DARPP-32 content (fig. S6A). (**D**) Digitized reconstruction of infected D2- and D1-SPNs in an entire DStr hemisection. (**E**) Percentage of particles classified as D2- and D1-SPNs (fig. S6B-D). (**F**) HSV1-H129Floxed virus (permanently switches from eGFP to tdTomato if encounters Cre) was bilaterally injected in the DStr of adora2a-Cre and drd1a-Cre mice. Infection in cis involves infection through isolated lineages (green and red). Infection in trans involves infection across lineages (yellow; see methods). (**H**) Viral spread and digitized reconstruction of red-labelled (tdTom) and yellow-labelled (tdTom + eGFP) SPNs in entire DStr hemisections of each Cre line. (**I**) Percentage of red and yellow SPNs in each Cre line (fig. S7). (**J**) Unilateral genetic lesion of D2-SPNs in the medial DStr (as in Fig. 2G). (**K**) Quantification of D2-SPN density (left axis) and taD1-SPN density (right axis) in control and lesioned sides after Rac + GBR pharmacological treatment (Fig. 3B). (L) Medio-lateral continuum from lesioned to intact DStr territories after drug treatment. Bottom: green (eGFP) and red (P-H3) fluorescence plot profile across the medio-lateral continuum above. M.G.V, mean grey value. *, significant overall/simple effect (black) and interaction (red). N.S., not significant (Table S4).

In order to confirm that the enhanced connectivity in D1-SPNs was influenced by a D2-to-D1 drive, we infected the dorsal striatum with HSV1 H129Floxed virus, an anterograde tracing approach in which transsynaptic labelling switches from green to red in the presence of Cre (27) (Fig. 4F; fig. S7, A to C). This method allowed us to quantify cross-system connectivity by assessing the proportion of (non Cre-expressing) neurons that received dual green and red infection within a critical temporal window (Fig. 4G, see Materials and Methods). Intra-striatal injection in adora2a-Cre mice revealed equal proportions of single (tdTom) and double (tdTom + eGFP) labeled SPNs, indicating that a large number of non Cre-expressing SPNs had received near simultaneous red and green viral influx (Fig. 4, H and I; table S4). By contrast, infection in drd1a-Cre mice produced a much lower proportion of double-labelled SPNs (Fig. 4, H and I; table S4), reflecting a reduced red and green influx in non Cre SPNs. Critically, the percentages of double and single-labelled neurons in drd1a-Cre mice matched those previously obtained with the non-switchable virus in drd2-eGFP (i.e. 20 and 80%, cf. Fig. 4, E and I). Again, these same proportions were consistently found across all infected striatal areas analyzed, irrespective of the area of spread (fig. S7, C to E). In sum, these results confirm that the two major projection systems in the striatum establish very different local connectivity, with D2-SPNs making substantial connections with D1-SPNs but not vice versa. The magnitude of this asymmetry goes well beyond that predicted by paired recordings and GABA inhibition experiments (28), and provides full neuroanatomical support for the large-scale D2-to-D1-SPN trans-modulation observed in this study.

To address the functional relevance of this connectivity bias, we evaluated the effects of RAC and GBR cocktail on striata where D2-SPNs were absent. We used the taCasp3-TEVp system (17) to induce genetic ablation of D2-SPNs in the DMS of one hemisphere (Fig. 4J). After recovery, these mice were treated with RAC and GBR prior to perfusion (as in Fig. 3B) and patterns of transcriptional activation were contrasted between control and lesioned sides. We found that the density of taD1-SPNs was inversely proportional to the density of D2-SPNs in control and lesioned sides (Fig. 4K; table S4). Moreover, analysis of P-H3+ nuclei across contiguous striatal territories spanning lesioned and intact areas revealed that the identity of the transcriptionally active neurons transitioned from D2-SPNs (in intact territories) to D1-SPNs (in D2-ablated territories) (Fig. 4L). These results confirm that activated D2-SPNs modulate molecular processes in neighboring D1-SPNs, and that they do so in situ through biased connectivity.

## D2-SPNs shape other sources of flexibility in goal-directed learning

Thus far, our results suggest that D2-SPNs shape the changes in striatal plasticity necessary for flexible encoding of goal-directed learning. In the case of inhibitory learning during extinction, this process is compatible with the role ascribed to DA in negative prediction error scenarios: a sustained pause of phasic DA in defined striatal territories can lead to recruitment of D2-SPNs and, with it, the D2-to-D1 trans-modulation reported here. We next sought to evaluate the role of D2-SPNs in flexible learning that should not overtly depend on a negative reward prediction error. To achieve this, we aimed to induce subtle changes in the identity predictions of pre-existing action-outcome relationships by reversing the outcome congruence between pairs of action-outcome associations. We trained mice with bilateral D2-SPN ablations in the DMS (Fig. 5A) and their lesion controls on two action-outcome associations (A1-O1 and A2-O2), which generated identical performance in both groups (Fig. 5B, table S5). We then verified whether both A-O contingencies had been correctly encoded by giving the mice an outcome-specific devaluation test, which evaluated the effect of sensory-specific satiety on one or the other outcome on choice between the two trained actions (29–31) (Fig. 5C). Both groups demonstrated that they correctly encoded the initial contingencies (i.e. A1-O1 and A2-O2); satiety on one of the outcomes (e.g., O1) reduced performance of the action associated with that outcome in training (A1; devalued) relative to the other, still valued, action (A2; valued) (Fig. 5C; table S5). We then explored whether these mice could incorporate new information by training them with the outcome identities reversed for 5 days (i.e., A1-O2 and A2-O1) (Fig. 5D; table S5) prior to a second outcome-specific devaluation test (Fig. 5E).

**Fig. 5.**
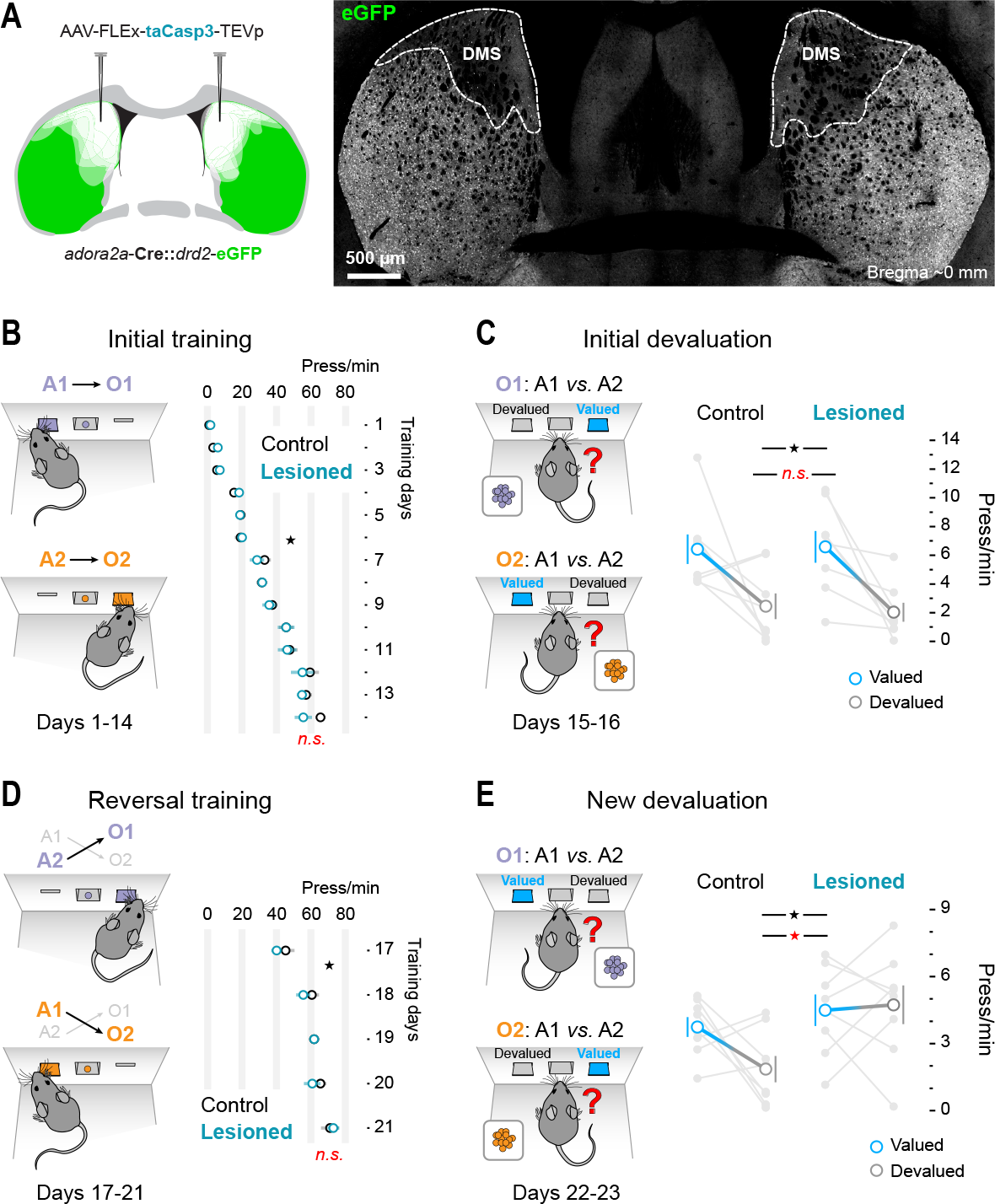
D2-SPNs control the addition of new learning. (**A**) Bilateral genetic lesion of D2-SPNs in the dorsomedial striatum (DMS) (Fig. 2G). Initial Learning: (**B**) Control and Lesioned mice were trained to two action – outcome (A-O) contingencies, resulting in elevated performance (press/min) across days. Initial devaluation test: a choice (A1 vs. A2) was presented after having sated the mice on one or the other outcome (O1/O2) over consecutive days. Graph shows performance on the valued (blue: provides non-sated O) and devalued (gray: provides sated O) levers. Additional Learning: (**D**) Mice were then trained to the reversed A-O contingencies, which rapidly elevated press/min performance. (**E**) A new round of devaluation and choice tests were presented (as in (C)). *, significant overall effect (black) and interaction (red). N.S., not significant (Table S5).

We found that, whereas Control mice were able to show flexible encoding and could adjust their choice according to the new A-O associations, mice with D2-SPN ablation failed to do so, as reflected by equal performance in Valued and Devalued levers on test (Fig. 5E; table S5). These results demonstrate that updating prior goal-directed learning critically involves intrastriatal postsynaptic D2-SPN function. Overall, we provide evidence of a ubiquitous D2-to-D1 trans-modulation mechanism supporting the integration of new with pre-existing learning in the striatum that goes beyond plasticity states induced by error-related phasic DA activity.

## Discussion

One of the most intriguing characteristics of the striatum is the random spatial distribution and high degree of intermingling between its D1-(direct) and D2-(indirect) projection systems, a feature that is actively promoted developmentally (32) and that has been meticulously retained throughout evolution (33). The result is a highly entropic binary mosaic that extends through an expansive and homogenous space and that is mostly devoid of histological boundaries (34). Such organization is unusual in the brain, and can be seen as an adaptation to provide an optimal postsynaptic scaffold for the integration of regionally meaningful neuromodulatory signals (35). In such a plain, borderless, environment, the rules established locally by D1 and D2-SPNs are likely to be critical in defining functional territoriality throughout the striatum, and this, we propose, is the key process shaping striatal-dependent learning.

Our study suggests that the striatum takes full advantage of this ‘one-to-one’ binary mosaic structure, in which activated D2-SPNs access and modify developing behavioral programs encoded by regionally defined ensembles of transcriptionally active D1-SPNs (what we call D2-to-D1 trans-modulation). We propose that this process is slow, as it depends on the molecular integration of additive neuromodulatory signals (7), but could, with time, create the regional functional boundaries that are necessary to identify and shape specific learning in the striatum. A good example of this sort of dynamic, persistent, neuromodulation is the recently described ‘wave-like’ motion of DA signals throughout the medio-lateral axis of the striatum (35). Beyond offering a broad solution to the credit assignment problem, recurrent waves of neuromodulatory activity in defined striatal areas could provide the kind of unbiased signal that, in the context of the molecular dichotomies established by D1 and D2 receptors (10), could shape the striatal mosaic into meaningful transcriptional motifs. In the case of extinction learning, as observed here, noisy alternations between DA rich and DA lean states within the DMS appears to generate a mixed population of activated SPNs comprising both D1 and D2 systems. This regional overlap lays the groundwork for the local ‘one-to-one’ modulation that shapes and integrates new learning, rewriting outdated D1-SPN function in the case of extinction learning, and segregating new and exerting territories of plasticity in the case of action-outcome identity reversal.

Our findings also suggest that the local bijective function of striatal mosaic modulation does not exclusively depend on DA prediction error-like signals, but likely encompasses other sources of local modulation, which are very abundant and heterogeneous in the striatum and that also have sharp dichotomic influence on SPN subtypes (36). One example—amongst many—is acetylcholine, which is locally sourced but tightly bound to DA (37), has an asymmetric influence over SPN subtypes (38) and has been shown to be critical for shaping goal-directed learning in the striatum (30, 31). Clearly, a more detailed understanding will be needed to determine the precise mechanisms that lead to the targeted recruitment of D2-SPNs to establish functional territoriality within the striatum. Acknowledging the collective and interdependent nature of the various neuromodulatory systems operating in the striatum, particularly those influencing transcriptional activity in SPNs, will provide a good starting point.

## Acknowledgements

We thank Z Skrbis for technical assistance.

