## Supplementary Materials for "D1 and D2 systems converge in the striatum to update goal-directed learning"

www.NeuroModuLab.org

**This PDF file includes:**

Materials and Methods
Figs. S1 to S7
Tables S1 to S5

### Materials and Methods

Animals

All experimental procedures were approved by the Animal Care and Ethics Committee at the University of New South Wales in accordance with the *Australian Code of Practice for the Care and Use of Animals for Scientific Purposes* (National Health and Medical Research Council, 2013). Adult (>3 months old) mice of both genders were used in these experiments. Behavioral experiments were performed on F1 outbred *drd2*-eGFP mice with C57BL6 background, whereas anatomy and pharmacology experiments were performed on inbred *drd2*-eGFP$^{+/+}$ mice. *Drd1a*-Cre$^{+/-}$ and *adora2a*-Cre$^{+/-}$ mice were used for the viral tracing experiments. In some experiments (see below), Cre$^{+/-}$ and eGFP$^{+/-}$ F1 hybrids (*drd1a*-Cre::*drd2*-eGFP and *adora2a*-Cre::*drd2*-eGFP) were obtained from inbred *drd1a*-Cre$^{+/+}$, *adora2a*-Cre$^{+/+}$ and *drd2*-eGFP$^{+/+}$ strains. Mice were housed in individually ventilated plastic home cages with a stand-alone air handling unit (Tecniplast, IT) in a climate-controlled colony room on a 12-h light/dark cycle.

Behavioral procedures and analysis

*Apparatus.* All training procedures were carried out in sound- and light-resistant modular operant chambers (Med Associates Inc, PO). The chambers were illuminated with a 3 W, 24 V house light. Each chamber contained a recessed feeding magazine at the center of the right side wall connected to an individual food dispenser that could deliver 20 mg Dustless Precision Pellets (Bio-Serv, NJ, #F0163) into the magazine when activated. Either side of the magazine contained retractable levers. Med-PC software was used to direct the insertion and retraction of the levers, illumination of the light and delivery of the pellets. The software allowed the recording of timestamp events each 10 msec, including the number of lever presses, magazine entries and food pellets delivered throughout the duration of the experiment. A combination of Med State Notation language and customized MATLAB (MathWorks, Natick, MA) scripts were used to retrieve timestamp data. Activity was also monitored using D-Link DCS-932L IP LED infrared cameras (VGA 1/5 inch CMOS sensor with built-in microphone) through D-ViewCam Software (D-link Corporation, Taiwan).

*Magazine training*. All mice weighted approximately 30 g at the beginning of each experiment, although food intake was restricted throughout the experiments (mice were maintained at approximately 85% of their free-feeding weight). Prior to all instrumental procedures, mice were exposed to four days of magazine training in the operant chambers (1 session/day). Each mouse was assigned one operant chamber, which was maintained throughout the duration of the experiment. The chamber light turned on to signal the beginning of the session and off at its termination. Both levers were retracted and mice explored the chamber freely. Twenty grain-based food pellets (20 mg, 3.35 kcal/g each) were delivered into the magazine at random time intervals (RT 60 seconds) over ~30 min. Consumption of pellets was monitored at the end of each session.

*Single-lever instrumental training.* These procedures were based on single-lever operant conditioning, where an action (lever press) was associated with the delivery of an outcome (food pellet). Mice were trained once daily. Each training session began when the test chamber light turned on and the lever contingent to the delivery of the pellets inserted in the chamber, and ended when the chamber light turned off and the lever retracted. Assignment of right or left lever was counterbalanced across mice. Each training session continued until mice had received a total of 20 pellets, or the session timed out at 30 min. For the first days of instrumental training, the lever delivered the pellet on a continuous reinforcement (CRF) schedule, in which every lever press was rewarded.

To study the effect of instrumental learning on striatal activation, one group (8 mice) was perfused early after establishing the instrumental contingency on the first day of CRF (Early group). Specifically, mice were perfused 20 min after reaching a criterion to detect the moment at which the A-O contingency was established. This criterion involved at least two first presses spaced by any interval (T=*n* sec), followed by two consecutive presses less than 60 sec apart (T<60 sec). This moment was stamped in the data as "first contingency" and triggered an additional "contingency time" counter, which was used to monitor the time post contingency in each mouse. Eight of the remaining animals (Late group) were trained to two additional days of CRF followed by a random ratio (RR) reinforcement schedule, in which the probability of the outcome given a response (P) was gradually shifted over 9 training sessions: RR5 (P = 0.2, 3 days), RR10 (P = 0.1, 3 days) and RR20 (P = 0.05, 3 days). The effect of learning on striatal activation was then assessed on the following day in an additional RR20 session (Late group). Finally, 8 yoked controls were also included (Yoked group), which received a reward whenever their 'contingent' littermate earned it through lever pressing. Therefore, these mice were exposed to the same conditions and food intake as their instrumental directors, but did not develop any action-outcome contingency as reward delivery was independent of their actions. Yoked animals were used to assess the specific effect of contingency on neuronal activity.

*Extinction learning and test.* In extinction of instrumental learning experiments, a more predictable fixed ratio (FR) reinforcement schedule was used, in which behavior was rewarded after a set number (N) of lever presses that increased over 9 training sessions: FR5 (N=5, 3 days) and FR10 (N=10, 6 days). On the following day, one group of 8 animals was maintained on the FR10 schedule (Instrumental group), whereas the remaining 8 animals were exposed to an instrumental extinction procedure, in which the previously trained response (lever press on FR10) was not reinforced (Extinction group). This procedure consisted of a pseudo-training session where all features of pellet delivery on an FR10 schedule were reproduced (i.e. vibration and sound of the pellet dispenser rotation), but no pellet was delivered. This allowed to maximize expectancy errors in mice and to monitor the times at which mice earned "norewards" (Ø). Instrumental and Extinction mice were perfused 20 min after the start of the session as described below. The same instrumental training was used in the experiment with 8 D2-SPN depleted (Lesioned) mice and 8 littermate controls (Unlesioned). However, after the last day of FR10 training, Lesioned and Unlesioned mice were exposed to a first 10-min extinction learning session and were placed back in their home cage. On the following day, mice were exposed to a 20-min extinction test, after which they were perfused as described below.

*Two-lever instrumental conditioning.* In reversal learning experiments, two action – outcome contingencies were trained in parallel (initial training). Each training session began with the illumination of the chamber light and insertion of one lever into the operant box, and ended with the extinguishing of the chamber light and lever retraction. The mice were exposed to two consecutive training sessions each day where only one lever was inserted into the box (and only the corresponding pellet delivered). The order of the sessions (purified or grain pellet first) was randomly alternated each day and lever-outcome assignment was fully counterbalanced. Pressing of the lever (right or left; A1 or A2) resulted in the delivery of the associated pellet (grain or purified; O1 or O2). Each training session (A1-O1 and A2-O2) continued until mice had received a total of 20 pellets, or the session timed out at 30 min. For the first three days of instrumental training (days 1-3), mice were trained on a constant reinforcement schedule (CRF); every lever pressing action was rewarded with the corresponding outcome. The probability of the outcome given a response (P) was gradually shifted over the following days of training using increasing random ratio (RR) schedules of reinforcement: a RR5 schedule (P = 0.2) was used on days 4–6, RR10 (P = 0.1) was used on days 7–9 and RR20 (P = 0.05) was used on days 10-14 and all further instrumental training (see below). While the probability of receiving a reward decreased over time, the initial contingencies remained unchanged (A1-O1 and A2-O2).

*Devaluation of outcomes.* On the last three days of initial instrumental training (days 12-14), mice were pre-habituated to devaluation boxes for 30 min to prevent novelty-induced stress on the days of devaluation. Pre-habituation was performed in a different room, where mice were placed individually into long, narrow boxes which contained an empty pellet container. Two devaluation trials were performed over two days (days 15-16), one outcome being devalued each day. The pellet container was filled with approximately 20 g of one or another outcome and animals were allowed to consume the reward to satiety over a 1 hr exposure period. Immediately following exposure, animals underwent an outcome-specific devaluation choice test in their assigned operant chambers.

*Outcome-specific devaluation choice test.* To test whether the animals developed a preference for the non-devalued outcome, a 10 min choice test was performed immediately after devaluation (Initial devaluation). Illumination of the chamber light and insertion of both levers indicated the beginning of the exercise. All testing was performed in extinction (pellet delivery did not occur, regardless of lever pressing). After 10 min, the light was extinguished and the levers were retracted, indicating the conclusion of the exercise. Two trials were conducted on consecutive days (one trial per outcome).

*Reversal of contingencies.* Further instrumental conditioning (RR20) was performed for five days following devaluation (days 17-21), with the initial contingencies now reversed (A1-O2, A2-O1, Reversal training). The precise combination of new lever-outcome assignments varied for each animal, depending on the animal's initially trained contingency. For the last two days of reversal training (days 20-21), the mice were pre-habituated for 30 min in the devaluation boxes (without food) prior to training. Following five days of contingency reversal training, the mice underwent two more days of outcome-specific devaluation choice test (days 22-23, New devaluation) (see above).

*Analysis of behavioral activity data.* Individual events (left or right lever press, magazine entries, and pellet delivery) that occurred during each training session were recorded. The time at which each event occurred was stamped with a 10 ms resolution. Datasets were imported into MATLAB and different approaches were used to analyze and visualize the data. To represent the automatization of instrumental actions throughout training we plotted return maps of identified behavioral elements (i.e. lever press and magazine check). Each map shows the performance of one mouse on the indicated day of training. Each dot corresponds to a lever press or magazine check (color coded), and includes the time delay to its preceding (x) and succeeding (y) element in the sequence. Data are Log10 transformed to approximate values. To analyze the effect of extinction on instrumental performance, we quantified lever presses occurring before each reward (or noreward) delivery across the session. Peri-event time histograms were used to visualize and quantify the collective lever press activity preceding each reward shown in chronological order.

Drug treatments

Raclopride (0.3 mg/kg; Tocris Cat. No. 1810) was dissolved in 0.9% (w/v) NaCl (saline) and GBR12783 dihydrochloride (15 mg/kg; Tocris Cat. No. 0513) was dissolved in water. The acute effect of these drugs on locomotor activity was measured in a square open field arena equipped with 3 sets of matching infrared photo beams that project across the open field along the three axes: X, Y, and Z (Med Associates Inc, PO). Photo beam breaks and the time at which they occurred were registered and data was processed using customised MATLAB scripts. Twenty-four homozygous *drd2*-eGFP mice were habituated to open field activity boxes and to intraperitoneal injections for 3 days. On the test day, animals were divided into four groups that received two injections (15 min apart) of different solutions: Vehicle-Vehicle; Raclopride-Vehicle; Vehicle-GBR12783; Raclopride-GBR12783. After each injection, mice were placed in the open field arena and 15 min after the second injection mice were perfused as described below. Locomotor activity (number of beam brakes) was measured at all times post injection, and behavioral trajectories were reconstructed from chronological beam brake data.

Viral procedures

*Inoculation of viruses.* Live viruses were injected into the brain through stereotaxic surgery. Mice were pre-anesthetized in an induction chamber with 5% isoflurane (Laser Animal Health, Pharmachem, Australia) delivered in oxygen (0.5L/min). Pedal reflex was used to monitor anesthesia before placing the mouse in the stereotaxic frame (Kopf Instruments) fitted with a mask supplying a continuous flow of oxygen/isoflurane mixture (0.5L/min). Prior to any incision, mice were given a subcutaneous injection of Carprofen analgesic (0.4 ml/kg). Once deeply anesthetized, a 1.5 cm incision was made to expose the skull. For each microinjection, a small hole was pierced (~0.2 mm width) at the appropriate A-P and M-L coordinates (see below). A pulled glass capillary (GC100TF-15, Harvard Apparatus) was pulled using a micropipette puller (P-97, Sutter Instrument) and pre-filled with the solution and fitted into a Nanoject III (Drummond Scientific) before being slowly inserted vertically through each hole until reaching target (D-V coordinate). Once in target, injections were preceded by a 2 to 5 min waiting interval. Different volume injections were delivered in one (unilateral experiments) or both (bilateral experiments) sides of the brain (see details below). Two to five min after the last pulse, the pipette was gently pulled out, and the incision was closed with surgery suture silk and sealed with tissue adhesive (3M Vetbond). Animals were given a s.c. injection of warm sterile saline (1mL) and placed in cages separately on a warming plate (37ºC) during recovery.

*Genetic ablation of D2-SPNs in adult mice.* Selective ablation of D2-SPNs was achieved through the taCasp – TEVp system (*17*). In brief, a designer procaspase 3 (taCasp3) lacking endogenous caspase cleavage sites but sensitive to the heterologous tobacco etch virus protease (TEVp) is expressed, together with the TEVp enzyme, in $Cre^+$ cells via AAV infection (AAV-Flex-taCasp3-TEVp; Addgene #45580). Twenty-one F1 hybrid *adora2a*-Cre::*drd2*-eGFP mice were unilaterally or bilaterally injected with Casp3-TEVp virus (Lesioned mice) or saline (Control mice) in the posterior dorsomedial striatum (A-P: +0.13 mm [bregma]; M-L: ±1.65 mm [bregma]; D-V: -2.6 mm [skull surface]). The injection (1 μL) was performed at a rate of 1 nL/sec over ~16 min. Pharmacological or behavioral procedures started after a 3-week recovery period. None of the mice showed perceptible wounds by the start of the experiments. The extent of the lesion was assessed by measuring the areas where *drd2*-eGFP had strongly diminished or cleared out.

*Intrastriatal collateral connectivity.* To assess the overall recurrent connectivity amongst striatal projection systems within the striatum, we expressed membrane-bound GFP (mGFP) and synaptophysin-mRuby (Syp-mRuby) in 3 F1 hybrid *adora2a*-Cre::*drd2*-eGFP and 3 *drd1a-Cre*::*drd2*-eGFP mice via AAV infection. This method allows to visualize anterograde synaptic territories of Cre expressing cells by labelling both axonal projections (mGFP) and pre-synaptic boutons (Syp-mRuby) (*24*). We injected the AAV5-hSyn-Flex-mGFP-2A-Syp-mRuby virus (Addgene #71760) bilaterally in the posterior dorsomedial striatum (A-P: +0.13 mm [bregma]; M-L: ±1.65 mm [bregma]; D-V: -2.6 mm [skull surface]). The injection (250 nL) was performed at a rate of 1 nl/sec over ~4 min. Mice were allowed to recover for 3 weeks prior to perfusion as described below.

*Transneuronal herpes viral tracing.* To perform transneuronal tracing, we employed fluorescent reporting (tdTomato) herpes simplex virus (HSV1), strain H129, which preferentially moves along synaptically connected neurons in the anterograde direction (*26*). In a first tracing experiment, $H129_{tdTomato}$ was injected unilaterally in the posterior dorsomedial striatum (A-P: 0.0 mm [bregma]; M-L: ±1.8 mm [bregma]; D-V: -2.85 mm [skull surface]) or the prelimbic cortex (A-P: 1.7 mm [bregma]; M-L: ±0.3 mm [bregma]; D-V: -2.1 mm [skull surface]) of 11 homozygous *drd2*-eGFP mice. The classification of infected particles into D2- ($eGFP^+$, DARPP-$32^+$) and D1- ($eGFP^-$, DARPP-$32^+$) SPNs allowed to calculate the proportion of each SPN subtype receiving anterograde transsynaptic infection, which informed about the extent of their overall connectivity. We allowed a 72-h incubation before perfusing the animals (as described below). During this period the virus is expected to spread no more than 2 or 3 synapses, therefore strongly representing both the local circuit network and the immediate extra-striatal circuitry. The experiment in the pre-limbic cortex was intended to inform about the influence of extra-striatal circuitry on the overall connectivity established in the striatum.

In a second tracing experiment, we employed a conditional neurotropic HSV1 (strain $H129_{Floxed}$) (*27*) that switches from expressing an eGFP (green) to a tdTomato (red) fluorescent reporter in the presence of Cre recombinase, and used this in combination with Cre transgenic mouse lines. With this approach, when a cell expresses Cre, the virus will drive the expression of tdTomato and thus initiate a red anterograde tracing lineage. On the other hand, in cells that do not express Cre, the virus will express eGFP and initiate a green anterograde lineage. Importantly, within a critical period, a Cre-negative cell receiving both red (from a Cre-expressing lineage) and green (from a non Cre expressing lineage) viruses may express both tdTomato and eGFP, and thus turn yellow. Quantification of the proportion of yellow-labelled and red-labelled neurons therefore allows estimations of the extent of cross-infection between striatal projection systems. $H129_{Floxed}$ was inoculated bilaterally in the posterior dorsomedial striatum (A-P: 0.0 mm [bregma]; M-L: ±1.8 mm [bregma]; D-V: -2.85 mm [dura]) of 4 *drd1a*-$Cre^{+/-}$ and 1 *drd1a*-$Cre^{-/-}$ or 4 *adora2a*-$Cre^{+/-}$ and 1 *adora2a*-$Cre^{-/-}$ mice to drive virus expression in D1- or D2-SPNs, respectively. Again, mice were allowed to recover for 72 h prior to perfusion, period in which the virus is expected to move along the local striatal and immediate extra-striatal circuits.

Tissue processing and immunofluorescence labelling

*Transcardial fixation and tissue sectioning.* While still conducting instrumental behaviours, mice were rapidly anesthetized by exposure to 10 sec of isoflurane gas in an induction chamber (4% in air; Laser Animal Health, Pharmachem, Australia), followed by a lethal intraperitoneal injection of sodium pentobarbital (0.5 mL of 10% Lethabarb; Virbac Pty. Ltd., Australia). After ensuring deep anaesthesia by testing paw and tail reflexes, mice were perfused transcardially using an air pressure system (constant flow of 15 ml/min) with cold formaldehyde solution (4%, w/v) in phosphate buffer saline (pH 7.4). Brains were dissected and post-fixed overnight in 4% formaldehyde solution at 4°C. Consecutive 30 µm coronal sections spanning the rostro-caudal extent of the striatum were obtained for each animal using a vibratome (VT1000s, Leica Microsystems, Germany). Free-floating slices were stored at -20°C in an anti-freeze cryoprotectant solution (ethylene glycol, 30% v/v; glycerol, 30% v/v, 0.25 M Tris buffer) until processed for immunofluorescence.

*Immunofluorescence staining.* Free floating brain slices were rinsed 3 times in Tris-buffered saline solution (TBS, 0.5 M) for 10 min on an orbital shaker at room temperature (R.T.) prior to membrane permeabilization treatment with triton X-100 solution (0.5%, v/v in TBS), which was applied for 2 h at R.T. After 3 10-min washes with TBS, slices were incubated at 4°C on an orbital shaker with rabbit polyclonal anti-phospho Histone H3 (Ser10) antibodies (P-H3; 1:500 dilution; Millipore, Cat. No. 06-570) and with mouse monoclonal anti-dopamine and cyclic adenosine monophosphate-regulated phosphoprotein, Mr 32 kDa antibodies (DARPP-32; 1:500 dilution; BD Biosciences, CA; Cat. No. 611520). Following a 48-h incubation period, unbound primary antibodies were washed off with 4 washes of TBS, and bound primary antibodies were detected through incubation for 1 h at room temperature with donkey anti-rabbit Alexa Fluor 546 (1:400 dilution; Thermo Fisher Scientific; Cat. No. A10040) and donkey anti-mouse Alexa Fluor 647 (1:400 dilution; Thermo Fisher Scientific; Cat. No. A31571) secondary antibodies. Unbound secondary antibodies were washed off through 2 10-min rinses in TBS and 2 10-min final washes in TB. Brain samples were then placed on Superfrost Plus coated slides (Thermo Fisher Scientific, MA) and mounted with a coverslip on Vectashield antifade mounting medium (Vector Laboratories, CA; Cat. No. H-1000). Slides were stored at 4°C in the dark until image acquisition.

Image acquisition and quantitative analysis

*Spinning disk confocal microscopy.* Fluorescent brain sections were imaged using a spinning disk confocal system equipped with the Diskovery multi-modal imaging platform (Andor Technology) and a Zyla 4.2 sCMOS camera (Andor Technology), that allows fast capture of large mosaic images and high-sensitivity high dynamic range imaging. The Diskovery platform was added to a Nikon Eclipse TiE microscope body with a motorized stage and the Nikon Perfect Focus System, and image acquisition was controlled by Nikon NIS-Elements software. Single high-magnification wide field-of-view images for each brain hemisphere were generated by automatically stitching multiple adjacent frames from within a defined area using the motorized stage. Individual images were manually rotated and aligned using Adobe Photoshop (Adobe Inc.), such that striata from different animals had approximately the same orientation.

*Nucleosomal response mapping.* High-resolution mosaics of brain samples obtained with NIS-Elements were quantitatively analyzed using Fiji software (*39*). The outer boundary of the striatum was manually delineated in each hemisection (freehand tool), the area of the region was measured ($mm^2$), and coordinates (x, y) of the points defining the line were obtained. To quantify the number of activated SPNs in *drd2*-eGFP$^{+/+}$ or +/- mice and retrieve their location across the striatum, images were thresholded based on P-H3 immunolabelling and individual nuclei were automatically detected using the "Analyze Particle" command in Fiji, which finds the edge of an object (i.e. nucleus), outlines its contour using the wand tool, creates an individual region of interest (ROI), and then determines the coordinates (x, y) of its center point (centroid). DARPP-32 staining was used to verify that only SPNs were included in the analysis. To determine whether an activated SPN was D1- or D2-type, the generated ROIs were overlaid on the eGFP image and the mean grey value (m.g.v) of all the pixels within the selection was then measured. Resulting neuronal data, including position (x, y coordinates), area and mean grey value of the eGFP channel for each detected ROI, were exported in a tab-delimited spreadsheet format and imported into MATLAB. Custom-made scripts were used to obtain the density of activated SPNs (P-H3 counts/$mm^2$) for each SPN population, based on an eGFP m.g.v threshold. The position of activated neurons within the tissue was digitally reconstructed by combining a line plot defined by the edge points of the striatum and a scatter plot with circles at the locations specified by the neurons' centroid coordinates, color-coded according to their eGFP m.g.v.

*Identification of activated neural ensembles.* To analyze the spatial distribution of activated D1- and D2-SPNs across the striatum and identify the extent of clustering, we used a density-based scan algorithm with noise (DBSCAN) approach written in MATLAB (*40*). The criteria used was that for each SPN belonging to a cluster, the neighborhood of a given radius (120 µm), needs to contain at least a minimum of 50 SPNs. To visually assess the density of activated neurons across striatal territories in each projection subsystem, we constructed distribution contours in MATLAB using the kde2d function, which estimates a bivariate kernel density over a set of grid points (*41*). To quantify how spatially intermingled activated D1- and D2-SPN clusters were in each analyzed striatum, we used Fiji to connect the outer points of up to 6 P-H3$^+$ D1- and D2-SPN ensembles delineating an ROI per each cluster, then measured the area and obtained x, y coordinates of the traced line. To quantify the degree of overlap between such ensembles, we calculated, for each image, the intersection areas between all D1- and D2-SPN cluster ROIs and obtained the x, y coordinates of the resulting outlines.

*Mapping of transynaptically labelled neurons.* A similar method described above to identify activated (P-H3$^+$) SPNs was also employed to quantify the proportion of tdTomato-labelled SPNs in *drd2*-eGFP mice inoculated with HSV-H129-tdTomato virus and *drd1a*-Cre or *adora2a*-Cre mice inoculated with HSV-H129-Floxed virus (see above). Individual ROIs for infected cells were found using the "Analyze Particle" command in Fiji on the basis of tdTomato labelling. These ROI sets were then overlaid on the DARPP-32 and eGFP images and fluorescent values for each channel from individual identified particles were obtained. Cells that were not DARPP-32 positive were excluded from analysis and a threshold eGFP value was used to classify SPNs into D1- or D2-type (HSV-H129-tdTomato experiment) or into Cre positive or negative (HSV-H129-Floxed experiment). Maps of infected SPN positions were reconstructed using MATLAB (see above) and dots were color-coded according to their classification.

### Statistical analyses

Statistical significance of the effects found in our data was assessed using IBS SPSS Statistics software (version 24; IBM Corporation, Somers, NY). The a priori alpha level was set at $p < .05$. To analyze the effect of one or more independent variables, Levene's test was first used to test the null hypothesis that the error variance of each dependent variable was equal across groups, and uni or multi-factorial ANOVA was then conducted. If one or more repeatedly measured variables were considered, uni- or multifactorial repeated measures ANOVA was conducted. Homogeneity of variance was tested through Mauchly's test of sphericity, and Greenhouse-Geisser corrections were considered when sphericity was not met. Mixed ANOVA analyses were conducted in experiments involving within-subject and between subjects factors. In instances where further statistical detail was appropriate, we conducted additional simple effects comparisons based on individual one-way ANOVAs or Bonferroni post-hoc comparisons. Statistical comparisons between two means were conducted through independent t-tests, and equal variances were assumed or not according to the Levene's test for equality of variances.

### Supplementary Figures

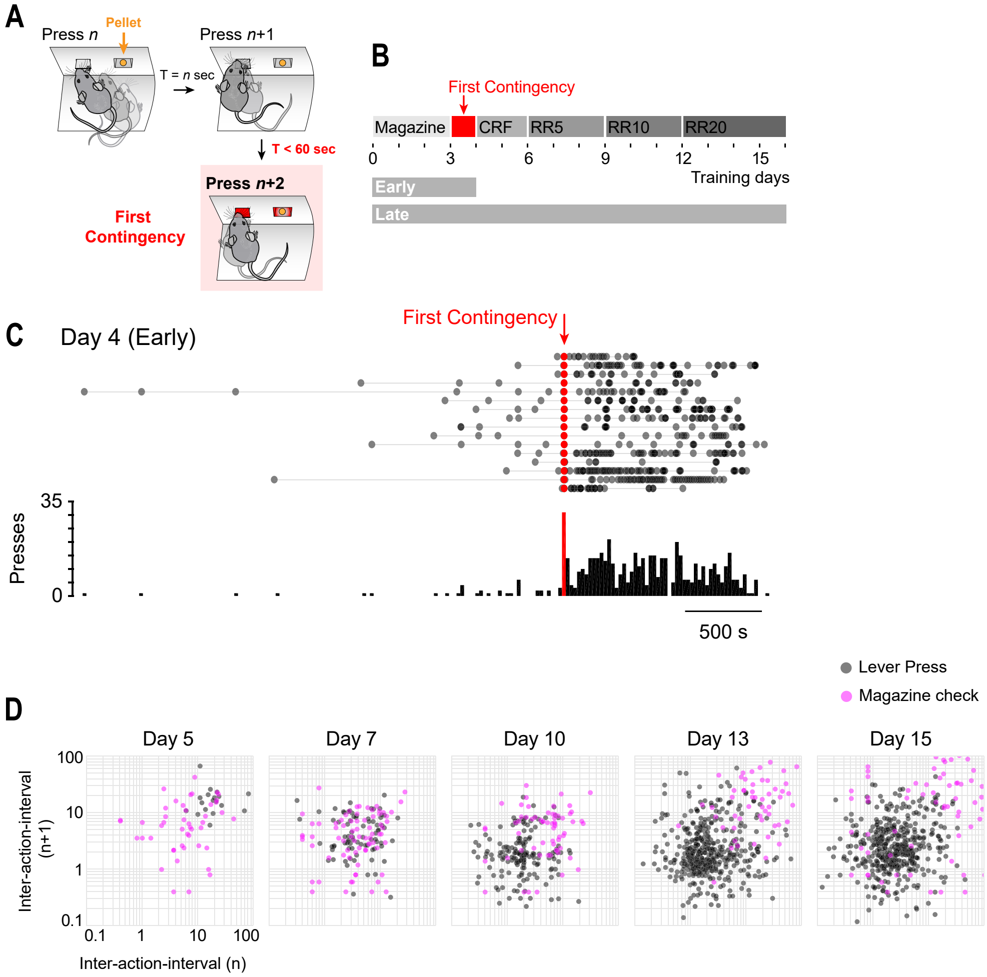


**Fig. S1. Early and late instrumental training produces characteristic patterns of lever press responding.** (**A**) A criterion for occurrence of the first association between an action (lever press) and an outcome (food pellet) was set to identify the approximate moment at which mice became spontaneously contingent. After the second press, any press that occurred within 60 seconds of the previous press triggered the "first contingency" signal. (**B**) Mice were trained to instrumental conditioning under an increasing random ratio (RR) schedule of reinforcement, from constant reinforcement (1 lever press = 1 outcome; CRF) to RR20 (~20 presses = 1 outcome). "Early" mice were perfused 20 min after the first contingency signal was triggered as in (A). "Late" mice were perfused 20 min after the start of RR20 on day 16. (**C**) Peri-event time histogram (PETH) of lever press activity aligned to the first contingency signal on day 4. (**D**) Instrumental signature developed by one example mouse (Group Late) across training. Plots are return maps of consecutive inter-action-intervals (IAI) on different training days showing how lever press patterns emerged as learning progressed. Each data point represents the time delay to its preceding (x) and succeeding (y) behavioral element. Grey: inter-press-intervals. Purple: inter-magazine check-intervals.

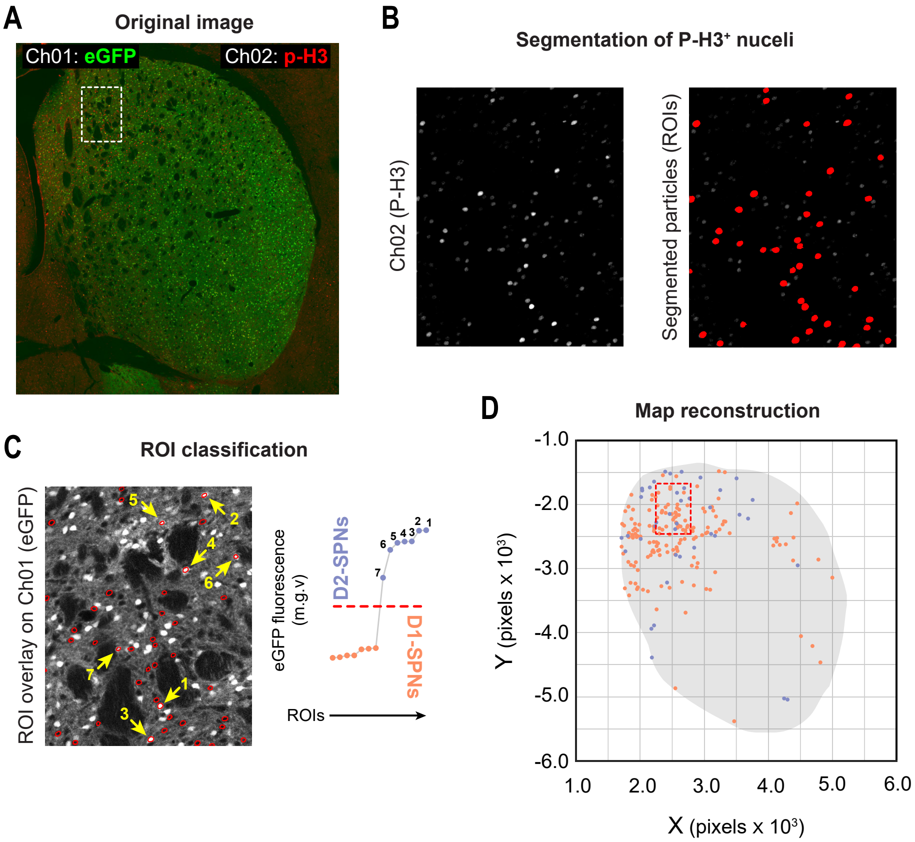


**Fig. S2. Nucleosomal response mapping in D2- and D1-SPNs of the dorsal striatum.** (**A**) Mapping was performed on high resolution (>6,000 px2) high bit depth (16 bit) confocal mosaics of superimposed *drd2*-eGFP (Ch01) and P-H3 (Ch02) signals. (**B**) Individual P-H3$^{+}$ nuclei (Ch02) were automatically detected through thresholding and segmentation methods, and regions of interest were defined around the edge of each particle. (**C**) Left: ROIs were superimposed on the eGFP image (Ch01), and the eGFP m.g.v. profile of each particle was obtained. Right: classification into D2- and D1-SPN was performed based on the bimodal distribution of eGFP m.g.v. profiles (high eGFP = D2-SPN; no eGFP = D1-SPN). Red dashed line indicates the division between distribution modes. (**D**) Digitized maps of each image were obtained by plotting the XY coordinate of the centroid of each classified particle (in pixels) in a bidimensional space.

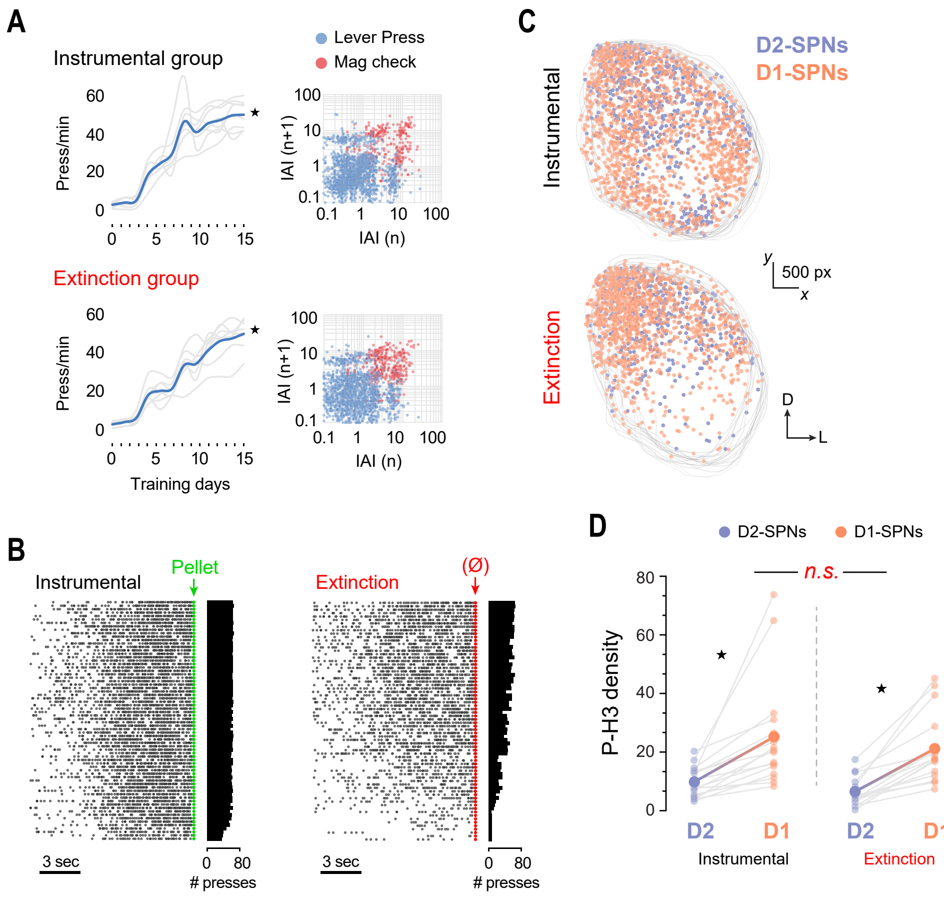


**Fig. S3. Execution and extinction of instrumental actions induce similar overall densities of P-H3$^{+}$ nuclei in D2- and D1-SPNs of the dorsal striatum.** (**A**) Average (blue trace) and individual (grey traces) lever press performance in both groups across instrumental training. Right panels show the return maps of IAIs displayed by each group on day 15 (pooled group data). (**B**) Raster plots and frequency histograms of pooled lever press activity data preceding the delivery of each reward (green; instrumental group) and pseudoreward (red; extinction group). (**C**) Digitized reconstruction of transcriptionally active (P-H3+) D2- and D1-SPNs throughout the dorsal striatum after 20 min of instrumental and extinction sessions (Figure 2A, day 16). Plots are superimposed individual maps. (**C**) Quantification of the P-H3$^{+}$ nuclei densities of each neuronal type in instrumental (top) and extinction (bottom) groups. Light color dots connected by light grey segments are paired D2- and D1-SPN densities in each image. * (black), simple effects; n.s., not significant (Table S2).

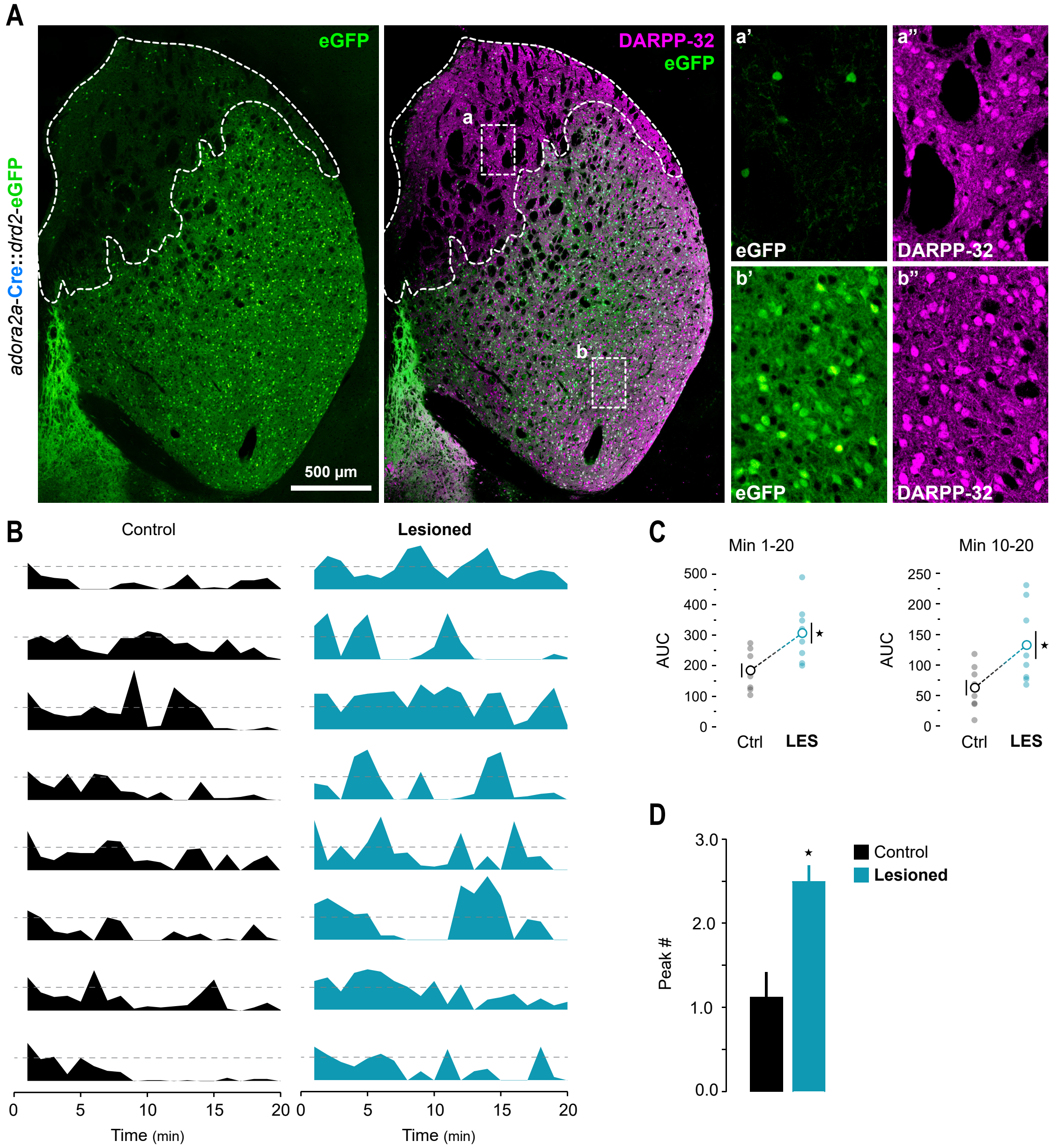


**Fig. S4. Genetic lesion of D2-SPNs in the dorsomedial striatum disrupts extinction learning.** Infection of AAV5-FLEx-taCasp3-TEVp virus results in Cre-dependent expression of the designer pro-caspase 3 (taCasp3) and tobacco etch virus protease (TEVp) (*17*). In the striatum of adult *adora2a*-Cre::*drd2*-eGFP transgenic mice (see Fig. 2G), TEVp cleaves and activates taCasp3 in neurons expressing A2A receptors, resulting in selective ablation of postsynaptic D2-SPNs. (**A**) Example of genetic lesion showing *drd2*-eGFP fluorescence (marks D2-SPNs) and DARPP-32 immunoreactivity (marks all SPNs) in the entire coronal dorsal striatum. The extent of the genetic lesion is evidenced by specific reduction of eGFP$^{+}$ neurons and/or eGFP$^{+}$ neuropil in the target region. Note how DARPP-32$^{+}$ GFP$^{-}$ SPNs are spared in lesioned territories. (**B**) Individual profiles of lever press activity per min in each mouse on extinction test (see Fig. 2H). Dashed grey lines mark the usual performance level at the start of the session (20 presses/min). Note how performance during extinction bounces back in lesioned but not control mice. (**C**) Quantification of the area under the curve (AUC) in the profiles in B (minute 1-20 and 10-20). (**D**) Quantification of the number of peaks above the 20 press/min mark. * (black), simple effects (Table S2).

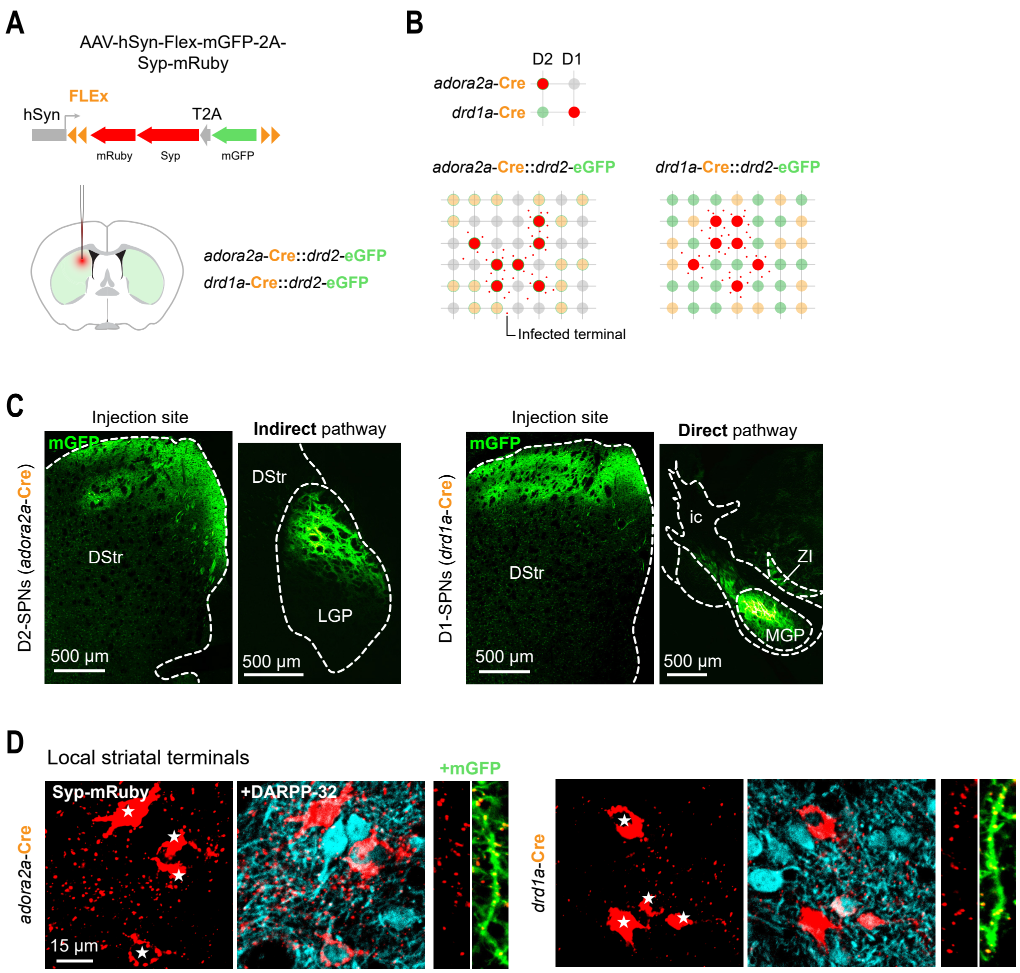


**Fig S5. D2- and D1- striatal projection systems display local presynaptic terminals in the striatum.** (**A**) Unilateral injection of AAV5-hSyn-FLEx-mGFP-2A-Synaptophysin-mRuby in the dorsal striatum (DStr). This virus allows tracing of specific cell types through Cre-dependent expression of mTagGFP (mGFP: marks axonal projections) and Synaptophysin-mRuby fusion protein (Sphyn-mRuby: marks presynaptic terminals) (24). (**B**) Expression of the virus in the striatum of *adora2a*-Cre::*drd2*-eGFP and *drd1a*-Cre::*drd2*-eGFP allows (1) visualizing local terminals derived from D2- and D1-SPNs, respectively, and (2) identifying subtype identity in surrounding SPNs. (**C**) This technique allows tracing of striatofugal pathways in the brain. Left: *adora2a*-Cre mediated expression of the virus in the DStr marks the indirect pathway as revealed by terminal staining in the lateral globus pallidus (LGP). Right: *drd1a*-Cre mediated expression in the same region marks the direct pathway, as revealed by terminal staining in the medial globus pallidus (MGP). (**D**) In the striatum, Sphyn-mRuby+ terminals were abundant in the striatal areas adjacent to the injection in both *adora2a*-Cre and *drd1a*-Cre mice. Although mRuby particles were observed close to mGFP$^{+}$ neurites of the same SPN type (insets), local terminals were widespread in the tissue and observed around the counter SPN type as well.

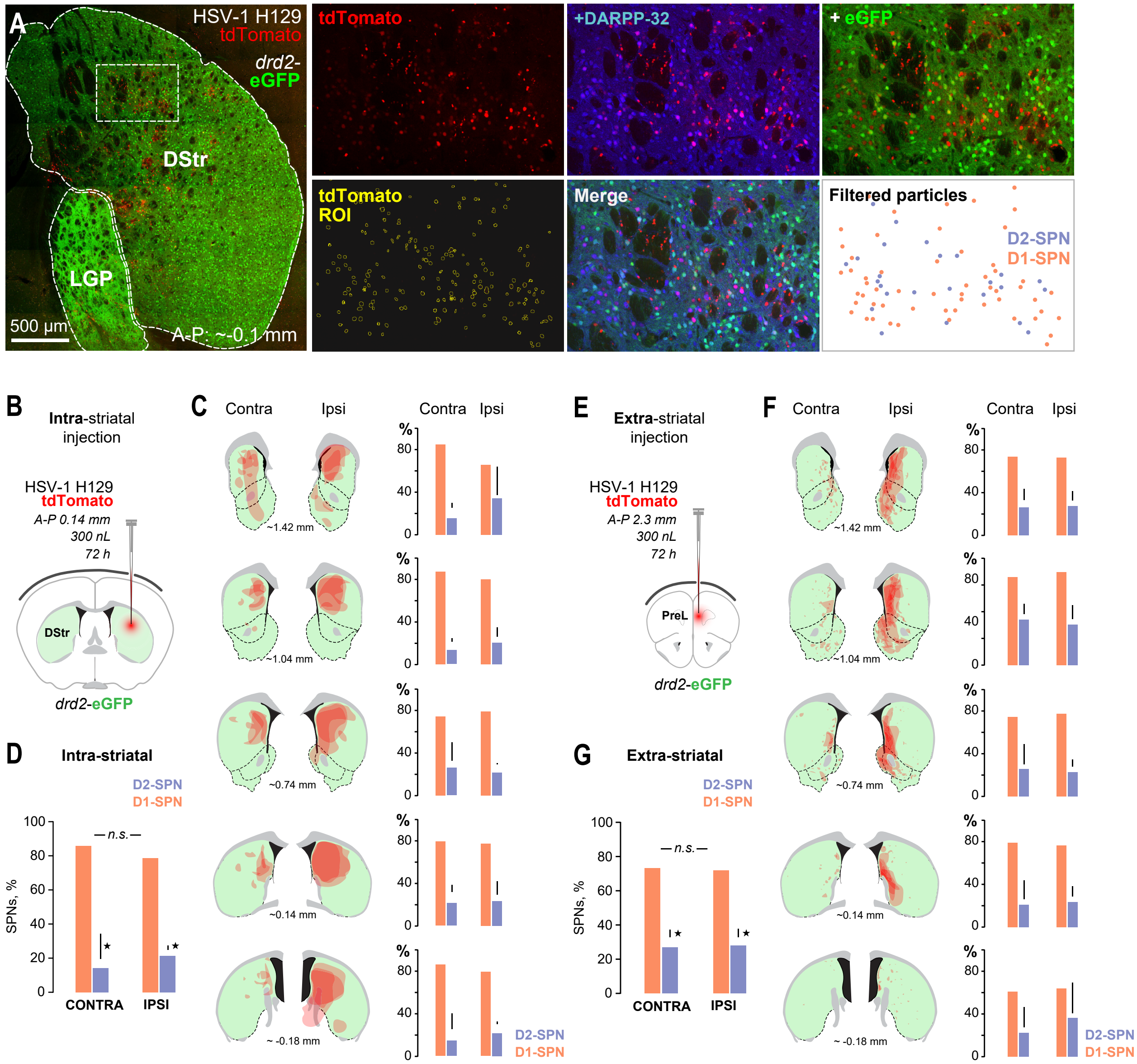


**Fig. S6. Striatal SPNs show a constant asymmetric pattern of trans-connectivity that is locally established in the striatum.** (**A**) SPN trans-connectivity quantification method. Cells infected with HSV1-H129-tdTomato anterograde trans-synaptic virus were identified in the entire coronal striatum at different rostro-caudal levels through semi-automated segmentation methods. Resulting particle ROIs were filtered according to DARPP-32 content (defines SPNs) and classified as D2- and D1-SPNs according to eGFP content. (**B**, **E**) The HSV-H129-tdTomato virus was unilaterally injected in the dorsal striatum (DStr) (intra-striatal injection; B) or the prelimbic (PrL) cortex (extra-striatal injection; E) and animals were perfused 72h later. (**C**, **F**) Diagrams showing the spread of tdTomato+ particles both ipsi- and contralateral to the injection at different rostro-caudal levels of the striatum. Right panels show the percentage of infected D2- and D1-SPNs corresponding to each level and side. A total of 3,673 tdTomato$^{+}$-SPNs were detected (131 SPNs/image on average) (**D**, **G**) Pooled percentage of infected D2- and D1-SPNs across the different rostro-caudal levels of the striatum. Distribution of SPNs was similarly disproportioned in both cases, irrespective of the injection site. * (black), simple effects (Table S4).

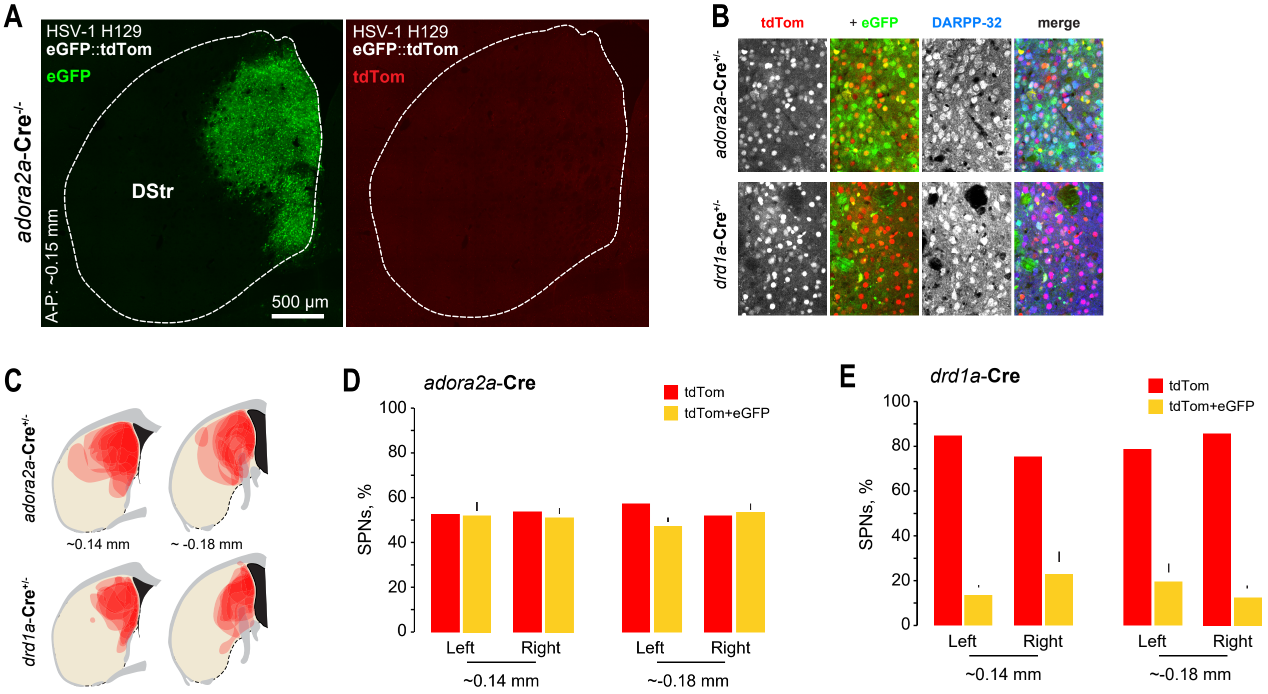


**Fig. S7. D2-SPNs show high trans-connectivity towards D1-SPNs.** (**A**) Expression of the non-switched HSV1-H129$_{\text{Floxed}}$ virus in the striatum of a control *adora2a*-Cre-/- mouse. (**B**) Representative images showing red-switched infection in *adora2a*-Cre and *drd1a*-Cre mice. Colocalization with DARPP-32 allowed filtering of particles corresponding to SPNs (similar to Figure S6A). Quantification of tdTomato$^+$ single labelled SPNs and tdTomato$^+$ + eGFP$^+$ double labelled SPNs informed about the amount of trans-connectivity from one system to the other (see Fig. 4G). (**C**) Diagrams showing the spread of the virus in the striatum at two different rostro-caudal levels in each transgenic line. (**D-E**) Percentage of tdTomato$^+$ single labelled and tdTomato$^+$ + eGFP$^+$ double labelled SPNs in *adora2a*-Cre (D) and *drd1a*-Cre (E) mice. Single labelled and double labelled particles reflect infection in *cis* and *trans*, respectively (see Fig. 4G). A total of 3,057 tdTomato$^+$-SPNs (in *drd1a*-Cre mice) and 3,274 (in *adora2a*-Cre mice) were detected.

**Table S1. Statistical Analysis 1 (Fig. 1 & S3).** Alpha level was set at $p < .05$. Significant effects are marked in bold.

| Fig. | Analysis | Test | Dep. variable | Factor(s) | Statistic | P value |
|---|---|---|---|---|---|---|
| 1D | Action rate (main effect) | One Way ANOVA | Action rate (press/min) | Training group (between subjects) | Main Effect: $F_{(2,21)} = 49,11$ | **< 0.001** |
| 1D | P-H3 density (main effect) | One Way ANOVA | P-H3 density (cells/mm$^2$) | Training group (between subjects) | Main Effect: $F_{(2,38)} = 17.29$ | **< 0.001** |
| 1D | Action rate (simple effects) | Bonferroni *post hoc* | Action rate (press/min) | — | Simple Effects:<br>Yoked vs. Early<br>Yoked vs. Late<br>Early vs. Late | <br>> 0.05<br>**< 0.001**<br>**< 0.001** |
| 1D | P-H3 density (simple effects) | Bonferroni *Post hoc* | P-H3 density (cells/mm$^2$) | — | Simple Effects:<br>Yoked vs. Early<br>Yoked vs. Late<br>Early vs. Late | <br>**< 0.001**<br>> 0.05<br>**< 0.05** |

**Table S2. Statistical Analysis 2 (Fig. 2, S3 & S4).** Alpha level was set at p < .05. Factor interactions are marked in red. Significant effects are marked in bold.

| Fig. | Analysis | Test | Dep. variable | Factor(s) | Statistic | P value |
|---|---|---|---|---|---|---|
| S3A | Instrumental acquisition | Two Way *mixed* ANOVA | Lever press rate (press/min) | (a) Training day (within-subjects)<br>(b) Group (between subjects) | Mauchly's sph. test: p < 0.001<br>Greenhouse Geisser correction:<br>Main Effect (training): $F_{(3.68,51.51)} = 191.73$<br>Training x Group interaction: $F_{(3.68,51.51)} = 2.56$ | **< 0.001**<br>> 0.05 |
| 2B | Extinction | Two Way *mixed* ANOVA | Cumulative lever presses | (a) Performance/t (within-subjects)<br>(b) Group (between subjects) | Main Effect (performance): $F_{(19, 266)} = 292.14$<br>Performance x Group interaction: $F_{(19, 266)} = 21.13$ | **< 0.001**<br>**< 0.001** |
| 2B | Extinction (overall effect) | Independent samples T-Test | Average lever presses | Group (between subjects) | $t_{(14)} = 3.45$ | **< 0.01** |
| S3D | P-H3 density | Two Way *mixed* ANOVA | P-H3 density (cells/mm$^2$) | (a) Neuron type (within-subjects)<br>(b) Group (between subjects) | Main Effect (Neuron type): $F_{(1,30)} = 41.94$<br>Neuron x Group interaction: $F_{(1,30)} = 0.27$ | **< 0.001**<br>> 0.5 |
| 2E | Overlap (% of D2) | Independent samples T-Test (equal variances not assumed) | % of taD2-SPN areas that overlap with taD1-SPN areas | Group (between subjects) | Levene's test: F = 6.28, p < 0.05<br>$t_{(23.41)} = -3.83$ | **< 0.01** |
| 2E | D2/D1 ratio | Independent samples T-Test | D2/D1 area ratio | Group (between subjects) | $t_{(30)} = 0.59$ | > 0.5 |
| 2F | Overlap (% of D1) | Independent samples T-Test | % of taD1-SPN areas that overlap with taD2-SPN areas | Group (between subjects) | $t_{(30)} = -2.63$ | **< 0.05** |
| 2F | D1/D2 ratio | Independent samples T-Test | D1/D2 area ratio | Group (between subjects) | $t_{(30)} = 0.03$ | > 0.5 |
| 2I | Instrumental acquisition | Two Way *mixed* ANOVA | Lever press rate (press/min) | (a) Training day (within-subjects)<br>(b) Group (between subjects) | Mauchly's sph. test: p < 0.001<br>Greenhouse Geisser correction:<br>Main Effect (training): $F_{(2.67,37.37)} = 98.78$<br>Training x Group interaction: $F_{(2.67,37.37)} = 0.53$ | **< 0.001**<br>> 0.1 |

| | | | | | | |
|---|---|---|---|---|---|---|
| 2J | Extinction learning | Two Way *mixed* ANOVA | Cumulative lever presses | (a) Performance /t (within-subjects)<br><br>(b) Group (between subjects) | Mauchly's sph. test: $p < 0.001$<br><br>Greenhouse Geisser correction:<br>Main Effect (performance): $F_{(1.48, 20.70)} = 95.54$<br><br>Performance x Group interaction: $F_{(1.48, 20.70)} = 0.95$ | **< 0.001**<br><br>> 0.1 |
| 2J | Extinction learning (overall effect) | Independent samples T-Test | Average lever presses | Group (between subjects) | $t_{(14)} = -1.83$ | > 0.05 |
| 2J | Extinction learning (slope analysis) | Linear regression slope comparison | Slope | Group (between subjects) | $F_{(1,16)} = 2.46$ | > 0.1 |
| 2K | Extinction | Two Way *mixed* ANOVA | Cumulative lever presses | (a) Performance/t (within-subjects)<br><br>(b) Group (between subjects) | Main Effect (performance): $F_{(19,266)} = 91.06$<br><br>Performance x Group interaction: $F_{(19, 266)} = 5.75$ | **< 0.001**<br><br>**< 0.001** |
| 2K | Extinction (overall effect) | Independent samples T-Test | Average lever presses | Group (between subjects) | $t_{(14)} = -3.28$ | **< 0.01** |
| 2K | Extinction (slope analysis) | Linear regression slope comparison | Slope | Group (between subjects) | $F_{(1,36)} = 122$ | **< 0.001** |
| S4C | Extinction (min 1-20) | Independent samples T-Test | AUC | Group (between subjects) | $t_{(14)} = -2.98$ | **< 0.05** |
| S4C | Extinction (min 10-20) | Independent samples T-Test | AUC | Group (between subjects) | Levene's test: $F = 5.33$, $p < 0.05$<br><br>$t_{(10.94)} = -2.664$ | **< 0.05** |
| S4D | Peak count (min 0-20) | Independent samples T-Test | Average number of peaks above 20 presses | Group (between subjects) | $t_{(14)} = -3.92$ | **< 0.01** |
| 2N | Overlap (% of D1) | Independent samples T-Test (equal variances not assumed) | % of taD2-SPN areas that overlap with taD1-SPN areas | Group (between subjects) | Levene's test: $F = 10.02$, $p < 0.05$<br><br>$t_{(21.06)} = -4.44$ | **< 0.001** |
| 2O | P-H3 density (DMS) | Two Way *mixed* ANOVA | P-H3 density (cells/mm$^2$) | (a) Neuron type (within-subjects)<br><br>(b) Group (between subjects) | Main Effect (Neuron type): $F_{(1,30)} = 101.8$<br><br>Neuron x Group interaction: $F_{(1,30)} = 11.61$ | **< 0.001**<br><br>**< 0.01** |

**Table S3. Statistical Analysis 3 (Fig. 3).** Alpha level was set at $p < .05$. Factor interactions are marked in red. Significant effects are marked in bold.

| Fig. | Analysis | Test | Dep. variable | Factor(s) | Statistic | P value |
|---|---|---|---|---|---|---|
| 3C | Ambulation (overall effect) | One Way ANOVA | Average number of beam breaks (min 15-30) | Treatment (between subjects) | $F_{(3,28)} = 15.56$ | **< 0.001** |
| 3C | Ambulation (simple effects) | Bonferroni *post hoc* | Average number of beam breaks (min 15-30) | — | Simple Effects:<br>Veh+Veh vs. Rac+Veh<br>Veh+Veh vs. Veh+GBR<br>Veh+Veh vs. Rac+GBR<br>Veh+GBR vs. Rac+GBR | <br>> 0.05<br>**< 0.01**<br>**< 0.05**<br>> 0.05 |
| 3D-G | Ambulation/t | Two Way *mixed* ANOVA | Number of beam breaks/2 min | (a) Breaks/t (within-subjects)<br><br>(b) Treatment (between subjects) | Mauchly's sph. test: $p < 0.001$<br><br>Greenhouse Geisser correction:<br>Main Effect (breaks/t):<br>$F_{(5.51,154.30)} = 5.514$<br><br>Performance x Group interaction:<br>$F_{(16.53,154.3)} = 5.64$ | <br><br><br>**< 0.001**<br><br><br><br>**< 0.001** |
| 3L | P-H3$^{+}$ nuclei | Two Way *mixed* ANOVA | Number of P-H3$^{+}$ nuclei | (a) Neuron type (within-subjects)<br><br>(b) Treatment (between subjects) | Main Effect (Neuron):<br>$F_{(1,40)} = 11.70$<br><br>Main Effect (Treatment):<br>$F_{(1,40)} = 306.41$<br>Treatment x Neuron interaction:<br>$F_{(3,40)} = 63.2$ | <br>**= 0.001**<br><br><br>**< 0.001**<br><br><br>**< 0.001** |

**Table S4. Statistical Analysis 4 (Fig. 4 & S6).** Alpha level was set at $p < .05$. Factor interactions are marked in red. Significant effects are marked in bold.

| Fig. | Analysis | Test | Dep. variable | Factor(s) | Statistic | P value |
|---|---|---|---|---|---|---|
| 4E | % of infected SPNs (overall effect) | Paired samples T-test | % of particles | Neuron type (within-subjects) | t(9) = 14.88 | **< 0.001** |
| S6D | % of infected SPNs (**intra-striatal** injection; ipsi- vs. contralateral to injection) | Two Way *repeated measures* ANOVA | % of particles | (a) Neuron type (within-subjects)<br>(b) Side relative to injection (within-subjects) | Main Effect (Neuron): $F_{(1,4)} = 323.89$<br>Neuron x Side interaction: $F_{(1,4)} = 2.49$ | **< 0.001**<br>> 0.1 |
| S6G | % of infected SPNs (**extra-striatal** injection; ipsi- vs. contralateral to injection) | Two Way *repeated measures* ANOVA | % of particles | (a) Neuron type (within-subjects)<br>(b) Side relative to injection (within-subjects) | Main Effect (Neuron): $F_{(1,4)} = 132.73$<br>Neuron x Side interaction: $F_{(1,4)} = 0.3$ | **< 0.001**<br>> 0.1 |
| 4I | % of infected SPNs (overall effect) | Two Way *mixed* ANOVA | % of particles | (a) Genotype (between subjects)<br>(b) Neuron type (within-subjects) | Main Effect (Neuron): $F_{(1,27)} = 211.52$<br>Genotype x Neuron interaction: $F_{(1,27)} = 177.42$ | **< 0.001**<br>**< 0.001** |
| 4I | % of infected SPNs (simple effects) | Paired samples T-test | % of particles | — | Simple Effects (tdTom+eGFP vs. tdTom):<br>*Adora2a*-Cre: $t_{(15)} = 1.07$<br>Drd1a-Cre: $t_{(12)} = 16.21$ | <br>> 0.1<br>**< 0.001** |
| 4K | Cell density (DMS) | Two Way *repeated measures* ANOVA | ROI density (ROIs/mm$^2$) | (a) Lesioned/control side (within-subjects)<br>(b) Neuron type (within-subjects) | Main Effect (Side): $F_{(1,4)} = 25.61$<br>Side x Neuron interaction: $F_{(1,4)} = 65.38$ | **< 0.01**<br>**< 0.01** |

**Table S5. Statistical Analysis 5 (Fig. 5).** Alpha level was set at p < .05. Factor interactions are marked in red. Significant effects are marked in bold.

| Fig. | Analysis | Test | Dep. variable | Factor(s) | Statistic | P value |
|---|---|---|---|---|---|---|
| 5B | Instrumental acquisition | Two Way *mixed* ANOVA | Lever press rate (press/min) | (a) Training day (within-subjects)<br><br>(b) Group (between subjects) | Mauchly's sph. test: p < 0.001<br><br>Greenhouse Geisser correction: Main Effect (training): $F_{(3.69,51.71)} = 83.84$<br><br>Training x Group interaction: $F_{(3.69,51.71)} = 2.56$ | **< 0.001**<br><br>> 0.1 |
| 5C | Initial devaluation (overall effect) | Three Way *mixed* ANOVA | Lever press rate (press/min) | (a) Lever (valued/devalued; within-subjects)<br><br>(b) Test (outcome 1/outcome 2; within-subjects)<br><br>(c) Group (lesioned/control; between subjects) | Main Effect (Lever): $F_{(1,14)} = 15.19$<br><br>Lever x Group interaction: $F_{(1,14)} = 0.66$<br><br>Main Effect (Test): $F_{(1,14)} = 16.21$<br><br>Test x Group interaction: $F_{(1,14)} = 0.074$ | **< 0.01**<br><br>> 0.5<br><br>< 0.01<br><br>> 0.5 |
| 5D | Reversal acquisition | Two Way *mixed* ANOVA | Lever press rate (press/min) | (a) Training day (within-subjects)<br><br>(b) Group (between subjects) | Mauchly's sph. test: p < 0.01<br><br>Greenhouse Geisser correction: Main Effect (training): $F_{(2.11,29.54)} = 22.82$<br><br>Training x Group interaction: $F_{(2.11,29.54)} = 0.55$ | **< 0.001**<br><br>> 0.5 |
| 5E | New devaluation (overall effect) | Three Way *mixed* ANOVA | Lever press rate (press/min) | (a) Lever (valued/devalued; within-subjects)<br><br>(b) Test (outcome 1/outcome 2; within-subjects)<br><br>(c) Group (lesioned/control; between subjects) | Main Effect (Lever): $F_{(1,14)} = 71.03$<br><br>Lever x Group interaction: $F_{(1,14)} = 5.17$<br><br>Main Effect (Test): $F_{(1,14)} = 1.68$<br><br>Test x Group interaction: $F_{(1,14)} = 2.81$ | **< 0.001**<br><br>**< 0.05**<br><br>> 0.1<br><br>> 0.1 |
